# XRN1 supplies free nucleotides to feed alphavirus replication

**DOI:** 10.1101/2024.12.09.625895

**Authors:** Vincenzo Ruscica, Louisa Iselin, Ryan Hull, Azman Embarc-Buh, Samyukta Narayanan, Natasha Palmalux, Namah Raut, Quan Gu, Honglin Chen, Marko Noerenberg, Zaydah R. de Laurent, Josmi Joseph, Michelle Noble, Catia Igreja, David L. Robertson, Joseph Hughes, Shabaz Mohammed, Vicent Pelechano, Ilan Davis, Alfredo Castello

## Abstract

Several RNA viruses induce widespread degradation of cellular mRNAs upon infection; however, the biological significance and mechanistic details of this phenomenon remain unknown. Here, we make use of a model alphavirus, Sindbis virus (SINV), to fill this knowledge gap. We found that SINV triggers cellular RNA decay through the exonuclease XRN1 and the 5’-to-3’ degradation machinery (5-3DM). These proteins accumulate at viral replication organelles (VROs) and interact with the non-structural protein 1 (nsP1), bringing mRNA degradation into proximity with vRNA synthesis. Our data suggest that monophosphate nucleotides released by cellular RNA decay are recycled through the salvage pathway to feed viral replications. Our work thus reveals a fundamental connection between cellular mRNA degradation and viral replication via nucleotides repurposing.

**Research highlights:**

- 5’-3’ RNA decay is essential for the replication of a wide range of viruses.
- XRN1 directly interacts with transcripts which are degraded during infection.
- RNA decay factors and salvage pathway members localise to viral factories.
- Supplying nucleosides to several 5-3DM deficient cells facilitates SINV infection.

## INTRODUCTION

The 5’-to-3’ degradation machinery (5-3DM) mediates the co-translational decay of cellular mRNAs. It consists of the exonuclease XRN1, decapping proteins DCP1 and DCP2, the decapping enhancer EDC4, and accessory co-factors, including PATL1 and DDX6. 5’-to-3’ RNA decay requires enzymatic removal of the 5’ cap by the decapping complex (Houseley and Tollervey, 2009; Łabno et al., 2016; Tuck et al., 2020). Degradation is then performed from the monophosphorylated 5’ end by the exonuclease XRN1, whose hydrolytic activity releases monophosphorylated nucleotides (Jinek et al., 2011; Pelechano et al., 2015). Where in the cell XRN1-mediated RNA degradation takes place has been subject to debate. Components of the 5-3DM accumulate in cellular membranelles organelles known as processing (P-)bodies (Borbolis et al., 2022; Brothers et al., 2023; Ripin et al., 2024). However, single molecule RNA analyses suggested that active degradation may occur in the cytosol with P-bodies serving as storage hubs for the 5-3DM and the translationally repressed RNAs (Blake et al., 2024; Brothers et al., 2023; Horvathova et al., 2017).

Beyond its key roles in maintaining cellular mRNA homeostasis, XRN1 has also been linked to virus infection, classed as dependency and antiviral factor. For example, flaviviruses harbour a pseudoknot in the 3’ untranslated region (UTR) of their genomes that stalls XRN1 during viral (v)RNA degradation, leading to the generation of subgenomic flavivirus RNA fragments (sfRNAs). sfRNAs are thought to inhibit the antiviral activity of type I interferons (IFNs) and reduce XRN1 activity, influencing flavivirus pathogenicity (Chapman et al., 2014a; Chapman et al., 2014b; Moon et al., 2012; Pijlman et al., 2008). XRN1 is an essential host factor for other positive sense single stranded (ss)RNA viruses, including Sindbis virus (SINV) and severe acute respiratory syndrome coronavirus 2 (SARS-CoV-2) (Garcia-Moreno et al., 2019; Grodzki et al., 2022). In SARS-CoV-2, the role of XRN1 appears to be mediated by a transcriptional feedback loop due to the widespread degradation of cytoplasmic mRNA initiated by RNaseL (Burke et al., 2022; Shehata et al., 2024). Cellular RNA degradation also occurs during the infection with DNA viruses such as gamma-herpesvirus (Abernathy et al., 2015) and in the negative sense influenza A virus (IAV), where the decay is triggered by a viral-encoded endonuclease (Khaperskyy et al., 2016). XRN1 was suggested to inhibit the activation of interferon beta in IAV (Liu et al., 2021) and the formation of double stranded (ds)RNA, preventing the activation of the protein kinase R (PKR) in vaccinia and measles viruses (BenDavid et al., 2022; Burgess and Mohr, 2015). Conversely, XRN1 has been proposed to be antiviral in vesicular stomatitis virus (VSV), a negative stranded virus causing degradation of vRNA (Laudenbach et al., 2021; Ng et al., 2020). XRN1 is also proposed to inhibit coxsackievirus and poliovirus infection through the autophagy-dependent mechanism (Delorme-Axford et al., 2018). Irrespective of whether XRN1 is a dependency factor, antiviral factor, or both, the molecular mechanisms underpinning its functions in infection remain poorly understood.

Here we present an alternative model, in which XRN1 and the 5-3DM drive mRNA degradation, providing free nucleotides to “feed” viral replication. This model highlights a fundamental and direct connection between cellular RNA degradation and viral replication, while still being compatible with the other known functions of XRN1.

## RESULTS

### XRN1 is an essential player in RNA virus infection

Our previous studies suggested that XRN1 is essential for SINV infection (Garcia-Moreno et al., 2019). To extend these findings, we tested SINV gene expression in HEK293 and A549 cells with partial (PKO) or full (KO) ablation of XRN1. Lack of XRN1 led to complete abrogation of viral capsid expression, which was consistent across cell clones and lines, while partial XRN1 depletion caused an intermediate phenotype reducing capsid levels by 70-50% (Fig.1a and S1a-c). We next performed single molecule fluorescence *in situ* hybridisation (smFISH) to assess XRN1 effects with quantitative and spatial resolution. We used SINV_nsP3-mScarlet_, which expresses a red fluorescence protein fused to the non-structural protein 3 (nsP3), to localise the viral replication organelles (VROs) (Kamel et al., 2024). SINV RNA and nsP3-mScarlet co-localised in the perinuclear area of wild type (WT) cells (Fig.1b). Neither SINV RNA nor nsP3 were detected in most XRN1 KO cells, whereas the few virus-positive cells exhibited residual amounts of viral molecules. An explanation for these results could be that viral particles fail to infect XRN1 KO cells. However, similar copy numbers of SINV RNA were detected in WT and XRN1 KO cells during early infection, based on smFISH and RT-qPCR-based quantification (Fig.1c and S1d), hinting at replication defects, rather than defective entry.

**Figure 1.**
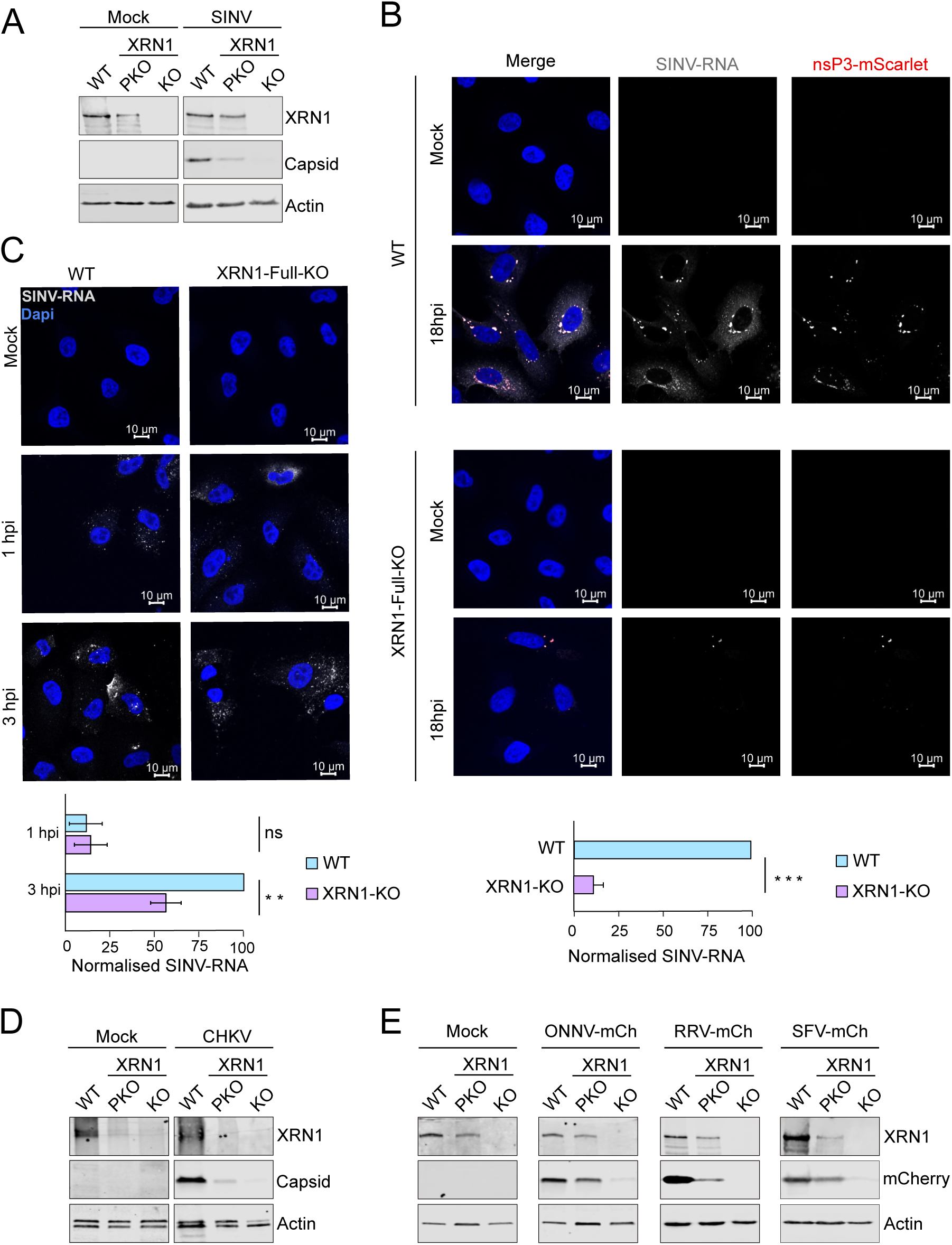
XRN1 is essential for alphavirus infection. (A) Western blot analysis of HEK293-WT, XRN1 partial (PKO) and full KO (KO) cells infected with SINV (MOI 0.1) for 18 h. n = 3. (B) Top: Fluorescence microscopy analysis of A549-WT and XRN1-KO cells infected with SINV_-nsP3-mScarlet_ for 18 h (MOI 0.5). vRNA was detected with smFISH using SINV specific probes, nsP3 was tagged with mScarlet, and nuclei were stained with DAPI. Bottom: Quantification of normalized SINV-RNA signal. n = 3; error bars: standard error; *** p < 0.001. (C) Top: Fluorescence microscopy analysis of A549-WT and XRN1-KO cells infected with SINV for 1 and 3 h (MOI 2). SINV RNA was detected with smFISH, and nuclei were stained with DAPI. Bottom: Quantification of normalized SINV-RNA signal. n = 3; error bars: standard error; ** p < 0.01. (D) Western blot analysis of HEK293-WT and XRN1 partial (PKO) and full KO (KO) cells infected with CHKV for 18 h (MOI 0.1). n = 3. (E) Western blot analysis of HEK293-WT and XRN1 partial (PKO) and full KO (KO) cells infected with ONNV-mCherry, RRV-mCherry, and SFV-mCherry for 18 h (MOI 0.1). n = 3.

We next explored the effects of XRN1 in the infection of other alphaviruses, including chikungunya virus (CHKV), O’nyong-nyong virus (ONNV), Ross River virus (RRV), and Semliki Forest virus (SFV). The levels of viral gene expression were strongly affected or completely abolished in XRN1 partial and full KO in all cases (Fig.1d-e). These results support that the central role of XRN1 is conserved across both old and new world alphaviruses.

We then extended our study to the positive sense ssRNA viruses coxsackievirus B3 (CVB3, *Picornaviridae*), Zika virus (ZIKV, *Flaviviridae*), human Coronavirus OC43 (OC43, *Coronaviridae*), severe acute respiratory syndrome coronavirus 2 (SARS-CoV-2, *Coronaviridae*), the negative sense ssRNA influenza A virus (IAV, *Orthomyxoviridae*), and the double stranded (ds)DNA vaccinia virus (VACV, *Poxviridae*). Lack of XRN1 caused an inhibition of CVB3, ZIKV, VACV, and SARS-CoV-2 infection ranging from strong to mild (Fig.S1e-h). Conversely, an increase in nucleocapsid accumulation was observed for the common cold coronavirus OC43, whose lifecycle is substantially slower compared to the other viruses tested (Fig.S1i) (Loo et al., 2020). Little to no effects were observed for IAV protein expression (Fig.S1j). Our results thus highlight the importance of XRN1 as dependency factor for a wide range of viruses belonging to different viral families, while also reporting cases in which XRN1 is not required (at least in the tested conditions). Moreover, the lack of phenotype in OC43 and IAV confirms that XRN1 KO cells do not have replication defects (Fig.S1k) and can sustain costly metabolic processes (Garcia-Moreno et al., 2019).

### Unveiling the critical roles of 5-3DM in virus infection

XRN1 harbours an exonuclease N-terminal domain, an intrinsically disordered region, and a C-terminal motif that mediates its interaction with EDC4 and recruitment of the decapping complex (Fig.S1a) (Chang et al., 2019). To test if the exonuclease activity and EDC4 interaction are important for sustaining viral replication, we expressed XRN1 mutants in which either of these molecular functions are disrupted. These mutants were expressed in a doxycycline-inducible manner in the XRN1 KO cells. Expression of XRN1-WT-Flag-HA caused a partial rescue of SINV gene expression, while a catalytic inactive mutant (D208A) (Jinek et al., 2011) failed to do so (Fig.2a). A mutant unable to bind EDC4 (Δ1650-1706) (Brothers et al., 2023; Chang et al., 2019) exhibited an intermediate phenotype (Fig.2a). Altogether, these data reveal that XRN1’s exonuclease activity is critical for SINV gene expression and its interaction with EDC4 is important to promote SINV replication.

**Figure 2.**
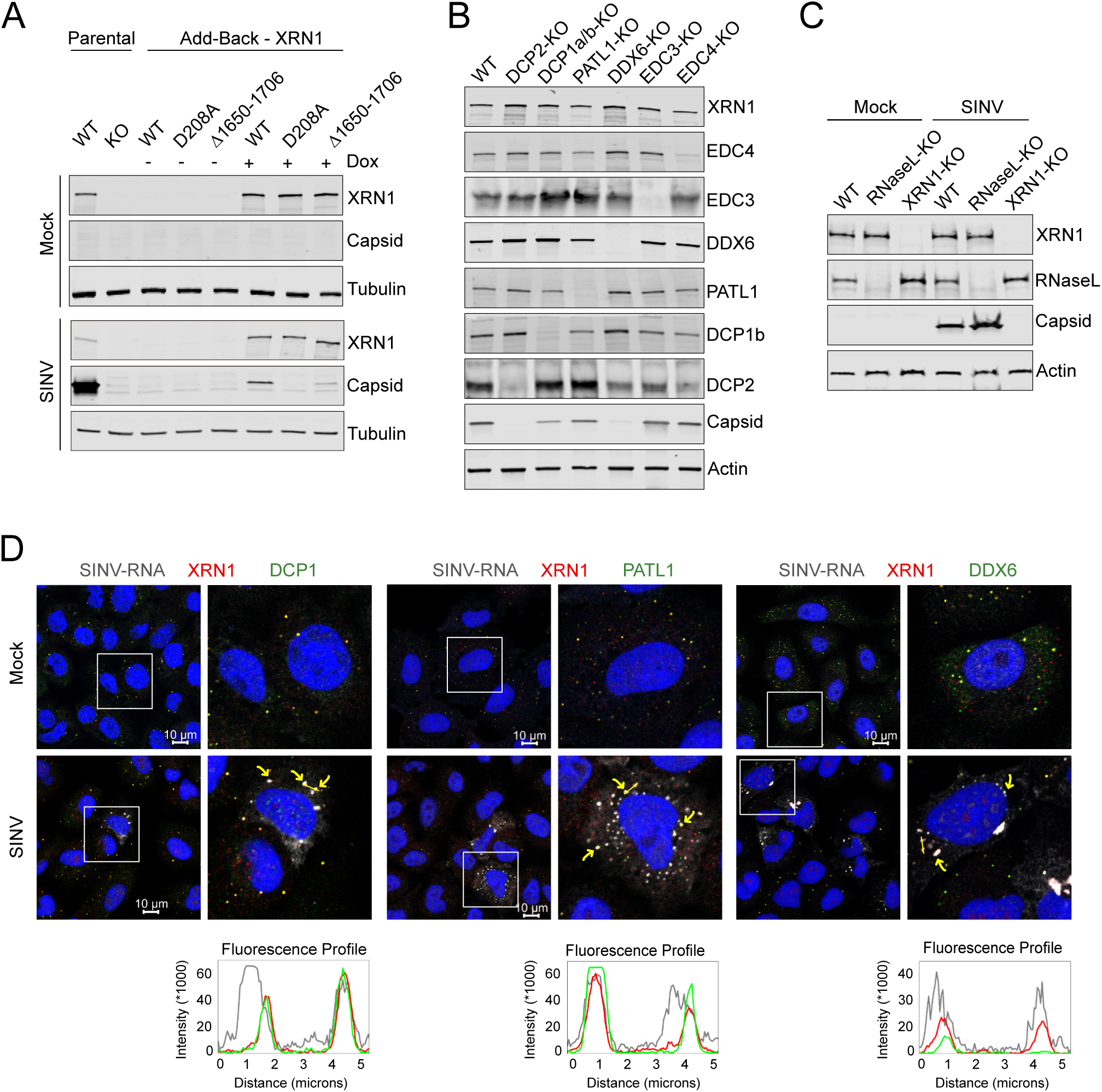
5’-3’ RNA decay pathway controls SINV infection. (A) Western blot analysis of HEK293-Flp-In T-REx WT and XRN1 KO cells complemented with the indicated XRN1 mutants and infected with SINV for 18 h (MOI 0.1). n = 3. Expression of XRN1 constructs, as shown in figure S1A, was induced by doxycycline treatment (Dox) for at least 96 h prior infection. (B) Western blot analysis of the indicated HEK293T-RNA decay KO cells infected with SINV for 18 h (MOI 0.01). n = 3. (C) Western blot analysis of HEK293 WT, RNaseL-KO, and XRN1-KO cells infected with SINV for 18 h (MOI 0.1). n = 3. (D) Fluorescence microscopy analysis of A549 WT cells infected with SINV for 18 h (MOI 0.5). n = 3. vRNA was detected with smFISH using SINV probes, nuclei were stained with DAPI, and XRN1, DCP1, PATL1, or DDX6 were visualized by immunofluorescence with respective antibodies. Yellow arrows indicate examples of colocalization between XRN1 and DCP1, PATL1, or DDX6 respectively, with SINV-RNA. Yellow lines mark the location of the fluorescence profile shown at the bottom. Fluorescence profiles of the indicated proteins and SINV-RNA intensity across 5 µm linear distance are shown below the microscopy images.

XRN1 WT-Flag-HA accumulated in puncta in the cytosol of uninfected cells, which are consistent with the known localisation of XRN1 in P-bodies (Fig.S2a). To assess the location of XRN1 WT and mutants during SINV infection we infected with a high multiplicity of infection (MOI) to achieve detectable level of viral gene expression even in mutant lines, where infection is strongly suppressed. XRN1-FLAG-HA foci were in close proximity to or overlapping with nsP3-mScarlet positive VROs (Fig.S2a). By contrast, the D208 catalytically inactive mutant formed larger puncta in the cytosol and exhibited poor co-localisation with nsP3-mScarlet (Fig.S2a). XRN1-Δ1650-1706 (EDC4-binding-null) mutant exhibited diffused nuclear and cytoplasmic distribution, failing to form puncta. However, there was a weak overlap with nsP3-mScarlet signal (Fig.S2a). These results suggest a correlation between the level of viral gene expression and the abundance of XRN1 in VROs.

XRN1 requires the generation of 5’ monophosphorylated ends on mRNAs that can be generated by decapping or endonuclease activity. We thus tested whether the decapping complex or the endonuclease RNaseL are required for XRN1 effects in infection. SINV gene expression was fully abrogated when the decapping enzyme DCP2 and DDX6 were ablated, whereas removal of other 5-3DM components such as DCP1, PATL1, and EDC4 caused partial effects (Fig.2b). Conversely, SINV gene expression was not reduced by RNaseL ablation (Fig.2c and S2b) (Li et al., 2016), stressing the importance of the decapping complex as the upstream factors of XRN1. Additionally, smFISH and immunofluorescence indicated that DCP1, PATL1, and DDX6 co-localised with XRN1 and SINV RNA at the VROs (Fig.2d), implying spatial proximity between the 5-3DM and the viral replication complex. SFV, CVB3, SARS-CoV-2, and IAV showed similar dependencies on the 5-3DM to SINV, while only DCP2 ablation slightly affected ZIKV infection. None of the KOs affected OC43, consistent with its XRN1 independence (Fig.S2c-h).

### SINV infection causes a widespread, XRN1-dependent degradation of cellular mRNAs

XRN1 may regulate viral gene expression by acting on cellular or vRNAs. In a previous study we observed a pervasive loss of cellular mRNAs in SINV-infected cells (Garcia-Moreno et al., 2019), and we hypothesised that XRN1 activity might be instrumental. Using smFISH with fluorescently labelled oligo(dT) probes, we observed that signal derived from cellular poly(A)+ RNAs decreases particularly in the cytosol of SINV-infected cells (Fig.3a). Conversely, no loss of cellular poly(A)+ RNA is observed in XRN1 KO cells, hinting at XRN1 as a causal effect of mRNA degradation in SINV-infected cells.

**Figure 3.**
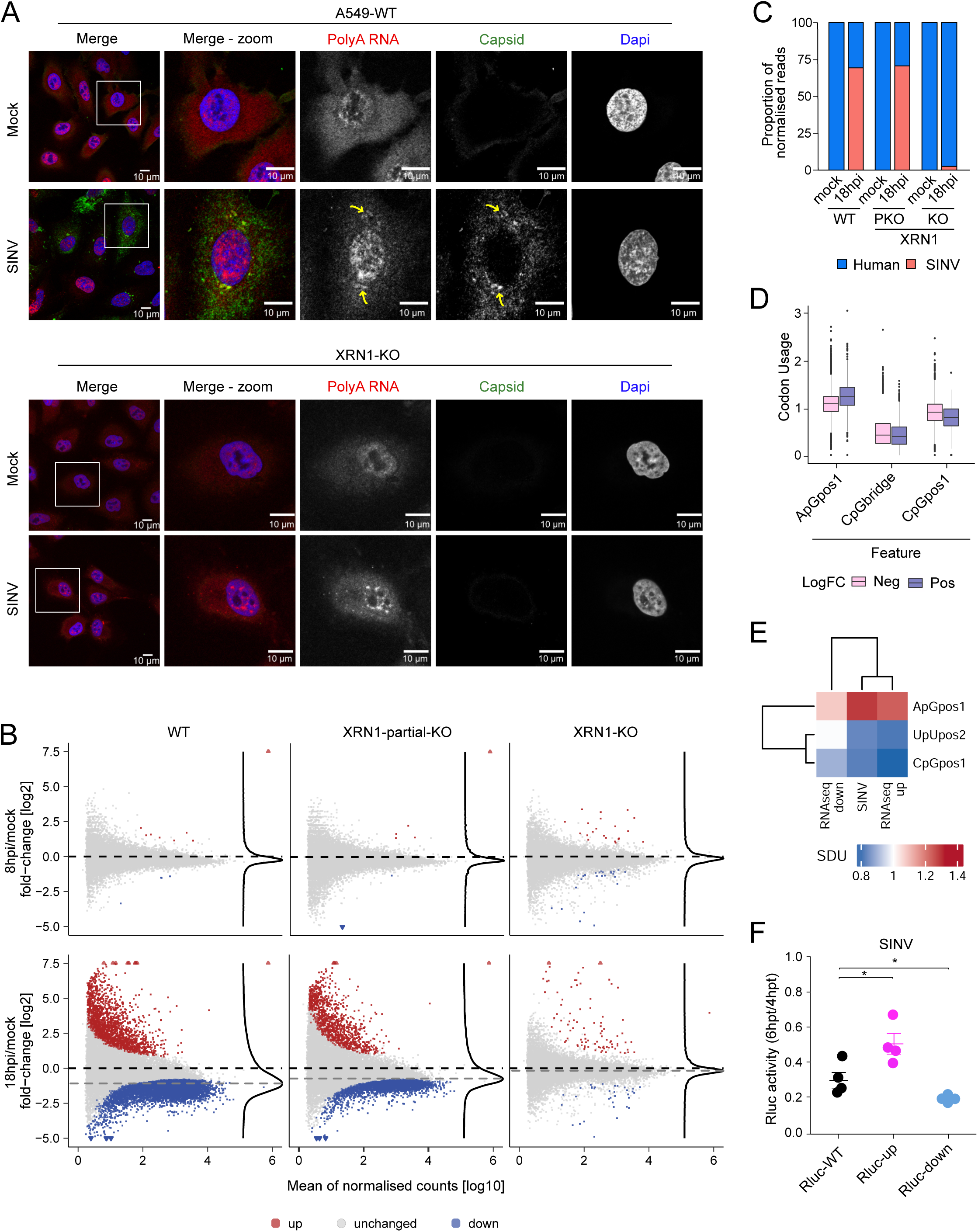
XRN1 induces widespread degradation of cellular mRNA. (A) Fluorescence microscopy analysis of A549 WT and XRN1-KO cells infected with SINV for 18 h (MOI 2). n = 3. Cellular mRNA was detected by smFISH with an Oligo(dT)_25_ probe (polyA RNA), nuclei were stained with DAPI, and SINV capsid was visualized by immunofluorescence. Yellow arrows indicate the accumulation of polyA RNA in the viral factories marked with SINV capsid antibody. (B) MA plot comparing the read coverage and the log2 fold change between SINV infected (8 hpi top; or 18 hpi bottom) and uninfected condition of each transcript detected in the RNA sequencing experiment for HEK293 WT, XRN1 partial, and full KO cells. Blue dots represent RNAs enriched with p.adj < 0.05 while grey dots represent non-significant changes. Dashed lines: black indicates the zero log2 fold change while grey represents the median fold changes across all transcripts. (C) Proportion of normalized human and SINV reads from RNAseq experiment at 18 hpi. (D) Boxplot of the Synonymous dinucleotide usage (SDU) of the three most important features in upregulated (pos) and downregulated (neg) transcripts upon SINV infection. (E) Comparison of the SDU of the indicated dinucleotides between SINV genome and cellular transcripts upregulated and downregulated upon SINV infection. (F) Luciferase activity of Rluc-WT or reporters with altered dinucleotide codon usage (Rluc-up and Rluc-down) measured at 6 hours post transfection (hpt) relative to the luciferase levels at 4 hpt in SINV infected conditions. n = 4; error bars: standard error; * p < 0.05.

To determine which RNAs are degraded in response to SINV, we employed RNA sequencing. Our results confirmed that SINV infection causes a pervasive degradation of housekeeping mRNAs, while vRNA and innate immunity genes increased in abundance (Fig.3b & S3a-c). Changes in cellular mRNA are still subtle at 8 hours post infection (hpi), but they become dramatic at 18hpi, implying that degradation occurs within the 8-18h timeframe (Fig.3b). Changes in abundance of cellular RNAs were confirmed by RT-qPCR (Fig.S3b). SINV RNA represents 70% of the poly(A)+ RNA at 18hpi, suggesting a biomass switch from cellular to vRNA (Fig.3c). An intermediate state in the RNA degradation landscape was observed in the partial KO, while viral RNA levels were comparable to those in WT cells (Fig.3b-c). Conversely, almost no effects in the cellular transcriptome were observed in infected XRN1 KO cells (Fig.3b-c), suggesting that cells failed to trigger global RNA decay despite being exposed to SINV.

XRN1 KO cells may be refractory to infection because of a higher basal level of interferon stimulated genes (ISGs) prior to infection, as suggested elsewhere (Burgess and Mohr, 2015; Liu et al., 2021; Zou et al., 2024). Conversely, we observed that most antiviral genes are expressed to a similar level in XRN1 KO and WT cells prior to infection (Fig.S3d). Indeed, no enrichment in GO terms related to innate immunity was observed in uninfected cells (Fig.S3e). Moreover, expression of the well-characterised ISG, OAS3, is similar in WT and KO lines. IFIT1 and OAS1 exhibited a moderate increase in few XRN1 KO clones. However, the deleterious effects of 5-3DM impairment in SINV infection were also observed in XRN1 KO clones with no effects in OAS1 and IFIT1 expression (Fig.S1b and S3f). Similarly, ablation of other components of the 5-3DM caused no alterations in OAS1 and IFIT1 expression (Fig.S3f). Altogether, these results strongly support that the inhibition of virus infection in these lines is not connected to a higher expression of ISGs.

The RNAseq data showed that a group of mRNAs escaped SINV-induced RNA degradation (Fig.3b). We compared the sequences of these two transcript populations to identify potential fate-determining features. We noticed differences in CpG and ApG dinucleotide frequency, particularly ApG in position 1 of upregulated transcripts and CpG and UpU in position 2 of downregulated transcripts (Fig.3d & S3g). Interestingly, SINV RNA had similar frequencies for these dinucleotides to the upregulated transcripts (Fig.3e and S3h). To assess if codon usage provided differential translation or stability features, we generated *Renilla* luciferase (Rluc) reporters with altered dinucleotides frequency while preserving amino acid frequency. Interestingly, the reporter containing the dinucleotide frequencies of SINV-resistant cellular mRNAs (Rluc-up) translates more efficiently in SINV infected cells, despite the virus induced shutoff of protein synthesis, than reporters with the frequencies of WT *Renilla* luciferase (Rluc-WT) or downregulated mRNAs (Rluc-down) (Fig.3f and S3i). While we have not detected differences in the stability of the reporter RNAs in the limited timeframe of the experiment (Fig.S3j-k), translation efficiency is directly linked with RNA decay (Bae and Coller, 2022; Morris et al., 2021; Presnyak et al., 2015; Weber and Chang, 2024).

### New insights into XRN1 mechanism of action from its RNA and protein interactomes

Our data highlighted the importance of XRN1 and the 5-3DM, but its connection to the RNA degradation in infected cells remains unclear. To test if XRN1 engages with the RNAs that are lost upon infection, we used iCLIP2 to generate a single-nucleotide resolution genome-wide map of XRN1 binding sites (Fig.S4a-b) (Buchbender et al., 2019). Our data showed that XRN1 increases its association with cellular mRNA at 4hpi, followed by a massive drop at 18hpi that correlates with the reduction of cellular mRNA substrates delineated by RNAseq (Fig.4a-b and Fig.3b). Analysis of XRN1 binding site distribution revealed that it accumulates at the 3’ end of target transcripts, particularly at 18hpi, when degradation is most pronounced. This is consistent with a slowdown of the exonuclease activity over coding regions, likely due to the higher incidence of secondary structure at the 3’UTRs (Fig.4c). By dividing RNAseq data from Fig3b based on whether genes are XRN1 targets or not, it became apparent that XRN1 preferentially associates with transcripts that are downregulated upon SINV infection (Fig.4d). Indeed, over 50% of XRN1’s targets are downregulated in WT cells at 18hpi, while a smaller fraction exhibit downregulation in PKO and almost none are downregulated in KO cells (Fig.4e). Furthermore, the iCLIP2 data shows evidence of XRN1 binding to at least 30% of the transcripts significantly downregulated during SINV infection at 4hpi (Fig.4f). This is an outstanding result given that iCLIP2 sensitivity is far from complete due to the limitations and biases of UV crosslinking and library preparation (Esteban-Serna et al., 2023; Hafner et al., 2021; Lee and Ule, 2018). XRN1 also binds to a large proportion of the mRNAs downregulated upon SINV infection in PKO cells, while this overlapping is not observed with the full KO (Fig.4f and Fig.S4c-d). These data highlight the physical interaction between XRN1 and the RNAs that are degraded in SINV-infected cells.

**Figure 4.**
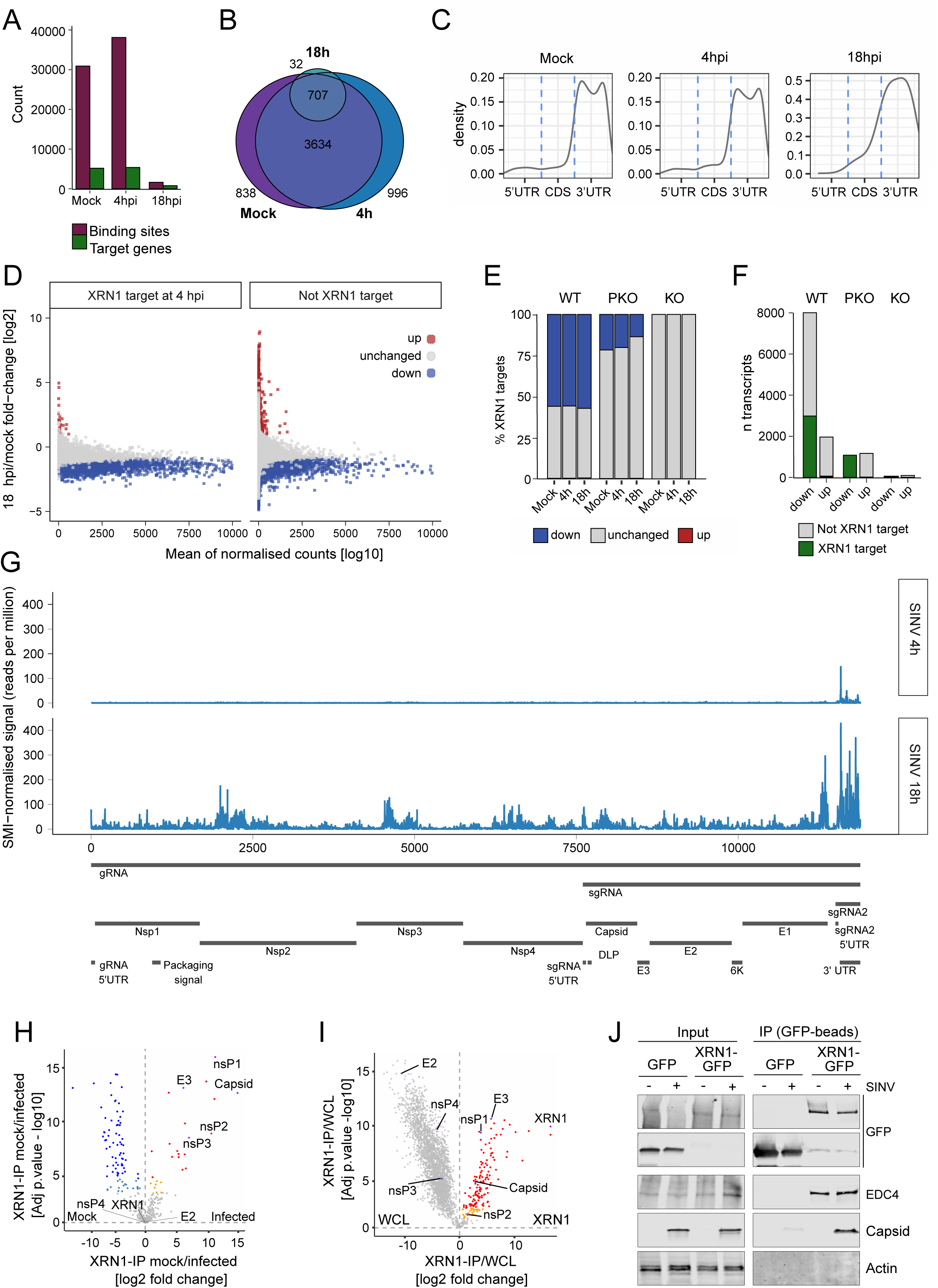
XRN1 directly binds RNAs which are degraded upon infection. (A) XRN1 binding site and target gene counts from iCLIP2 analysis of HEK293-Flp-In T-REx-XRN1-eGFP cells. Cells were either uninfected or infected with SINV for 4 or 18 h (MOI 10). n = 3. (B) Venn diagram representing the overlap of target genes identified in iCLIP2 in mock and SINV infected cells (4 h and 18 h). (C) Density plot of distribution of XRN1 binding sites on mature target mRNAs. (D) MA plots from figure 3B (of WT 18 hpi sample) showing XRN1 targets (left, 4hpi) and non-targets (right) identified by iCLIP2. Blue and red dots represent significantly downregulated and upregulated RNAs, respectively. (E) Bar plots showing the percentage of XRN1 target genes (based on iCLIP2) that are downregulated, unchanged, and upregulated (based on RNAseq) in WT, XRN1 partial (PKO), and full KO (KO). (F) Bar plots showing the number of XRN1 target genes and non-target genes (based on iCLIP2) that are upregulated and downregulated (based on RNAseq) in WT, XRN1 partial (PKO), and full KO (KO). (G) Top: Binding profile of XRN1 on SINV genome at 4 hpi and 18 hpi. Bottom: schematic representation of SINV genome features. (H) Volcano plot showing proteins enriched in HEK293-Flp-In T-REx-XRN1-eGFP IP in mock versus SINV-infected cells (18 hpi, MOI 10). n = 3. (I) Volcano plot showing proteins enriched in HEK293-Flp-In T-REx-XRN1-eGFP IP compared to WCL in SINV-infected cells (18 hpi, MOI 10). n = 3. (J) Western blot analysis of input and anti-GFP-beads IP (pull down of GFP-CTRL and XRN1-GFP) in mock and infected conditions. n = 3.

An alternative reason for the effects of XRN1 KO could relate to the generation of degradation-dependent sgRNAs as with flaviviruses (Slonchak and Khromykh, 2018). To confirm these results, we examined the iCLIP2 reads mapping to SINV RNA. XRN1 binding to vRNA was almost negligible at 4hpi, although it augmented at 18hpi (Fig.4g). This is expected as levels of vRNA are far superior at the late stages of infection. However, XRN1 binding sites on vRNA over cellular RNAs were substantially lower compared to RBPs known to interact with SINV such as TRIM25 or GEMIN5 (Fig.S4e) (Álvarez et al., 2024; Garcia-Moreno et al., 2019). These results suggest that although vRNA is not the primary target of XRN1, degradation of vRNA is likely to occur to some extent. The binding profile showed an increase in binding in the 3’ UTR, which is consistent with our previous observations for cellular RNAs (Fig.4c). To test if this increase in local binding could contribute to the formation of sgRNAs, we analysed the total RNAseq focusing on the regions downstream the most abundant binding sites. We did not observe differences in read density across the viral genome in WT and XRN1 KO lines, making it unlikely that XRN1-dependent SINV sgRNAs exist (Fig.S4f). Hence, we could not find evidence supporting the formation of XRN1-dependent sgRNAs in SINV infected cells.

The location of XRN1 and the 5-3DM proximal to VROs (Fig.2d) suggests a connection between cellular RNA decay and viral replication. To determine if there is a viral protein that tethers XRN1 to the viral factories, we performed affinity purification of XRN1-GFP in infected and uninfected cells followed by proteomics (Fig.S4g). As expected, we detected the 5-3DM complex in the eluates of both infected and uninfected cells (Fig.S4h-i). Additionally, we observed that XRN1 interacted with three proteins of the SINV RdRp complex: the capping pore nsP1, the helicase/protease nsP2, and nsP3 which is an interaction platform projected outside the VROs (Fig.4h) (Kamel et al., 2024; Kril et al., 2024; Tan et al., 2022). We did not detect RdRp nsP4, which is present in the inner side of the VROs and is substoichiometric to other components of the replication complex (Jones et al., 2021). Viral proteins were absent in mock controls; therefore, we compared the eluates against the whole cell proteome to exclude enrichment because of artefactual intensity ratios. This analysis confirmed a strong enrichment of nsP1, nsP2, capsid, and E3 in XRN1 IP eluates over the whole proteome, with nsP1 being the most enriched protein from the RdRp complex (Fig.4i). This is exciting as nsP1 multimerises to form a pore at the membrane of the VROs that controls metabolite trafficking, and vRNA capping and exiting from the VROs (Jones et al., 2021; Jones et al., 2023; Laurent et al., 2022; Tan et al., 2022; Zhang et al., 2022). XRN1-GFP IP followed by Western blotting with specific antibodies orthogonally confirmed our proteomic results (Fig.4j).

### Importance of the nucleotide salvage pathway in XRN1 proviral function

The role of XRN1 in infection thus far remains unknown. An exciting possibility is that XRN1-dependent degradation of cellular RNA supplies free nucleotides to the viral RdRp. However, XRN1 degradation releases mono-phosphorylated nucleotides (Chang et al., 2011; Jinek et al., 2011), while the RdRp requires tri-phosphorylated nucleotides (Chandel, 2021). If this hypothesis is correct, we expect to find the members of the nucleotide salvage pathway in proximity with the VROs. Immunofluorescence analysis revealed that, as hypothesised, nucleotide-modifying enzymes, particularly NME3, HPRT1, and APRT, partially co-localise with VROs (Fig.5a & S5a). We further investigated the involvement of the salvage pathway by simultaneously depleting APRT, HPRT1, UPRT, CTPS1/2, CDA, and NME3 via siRNA pools, and observed that their partial depletion leads to decreased viral replication without affecting cell viability (Fig.5b and S5b). Altogether, these results point to the salvage pathway as a facilitator of SINV replication.

**Figure 5.**
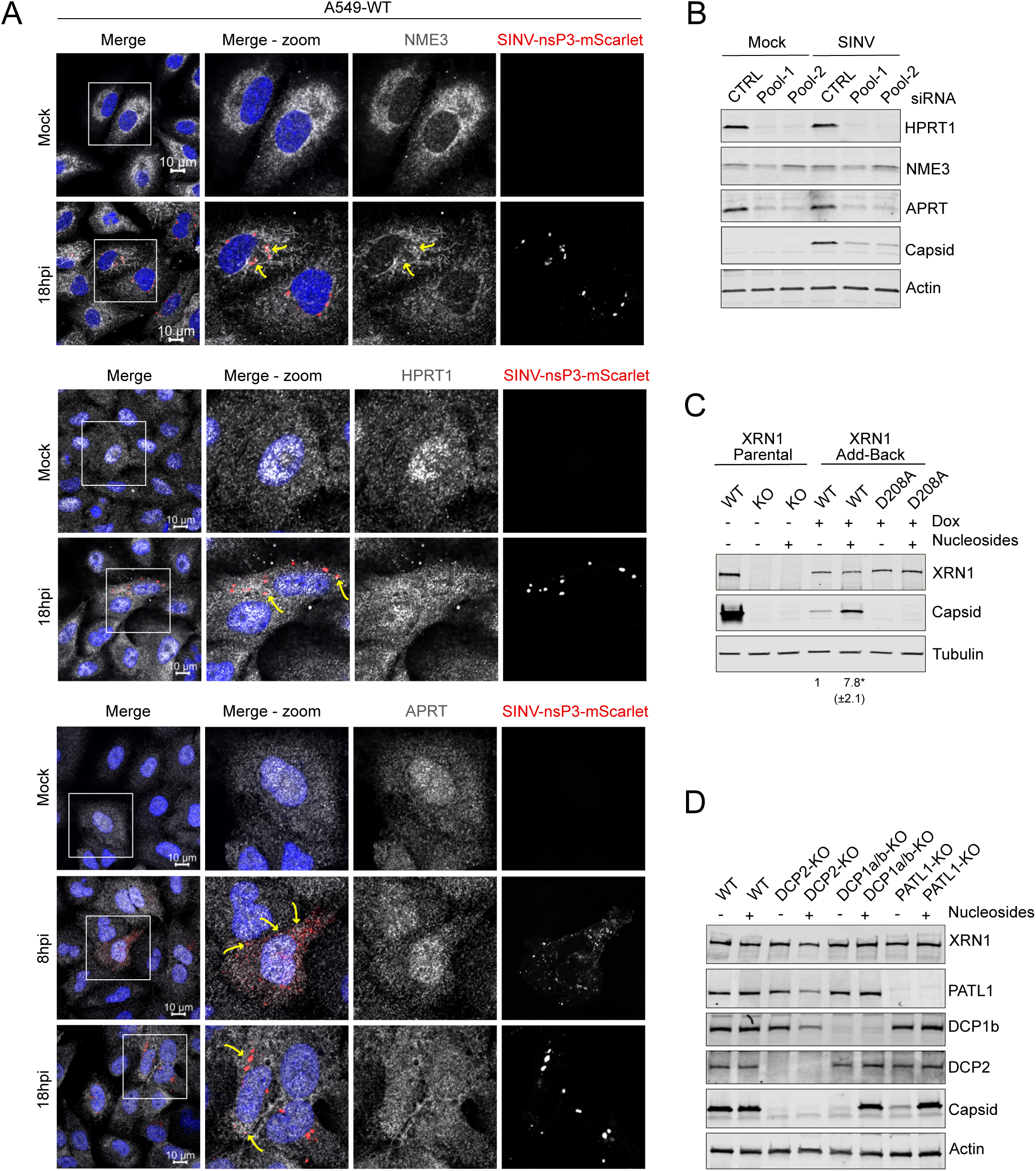
Providing nucleosides facilitates SINV infection. (A) Fluorescence microscopy analysis of mock and SINV-infected cells at the indicated time-points post infection (MOI 1). N = 3. NME3, HPRT1, and APRT were detected by immunofluorescence, SINV-nsP3 was visualized by tagging with mScarlet, and nuclei were stained with DAPI. Yellow arrows indicate colocalization of nsP3 with NME3 (top) HPRT1 (middle), and APRT (bottom). (B) Western blot analysis of A549-WT cells (CTRL siRNA) and cells depleted of components of the nucleotide salvage pathway with two different siRNAs mix (Pool-1: HPRT1, CTPS1/2, APRT, UPRT, CDA, and NME3; and Pool-2: HPRT1, CTPS1, APRT, CDA, and UPRT), infected with SINV for 18 h (MOI 0.1). n = 3. (C) Western blot analysis of HEK293-Flp-In T-REx WT and XRN1 KO cells complemented with the indicated XRN1 mutants, in presence or absence of externally supplemented nucleosides and infected with SINV for 18 h (MOI 0.1). n = 3. Below the blots the average normalized capsid level of three independent experiments, in bracket standard error. * p < 0.05. Expression of XRN1 constructs was induced by doxycycline treatment (Dox) for at least 96 hours prior infection. (D) Western blot analysis of HEK293T KO cells of the indicated member of the RNA decay machinery, supplemented with nucleosides (2X) and infected with SINV for 18 h (MOI 0.01). n = 3.

If XRN1 roles in infection are associated with nucleotides supply, we reasoned that external supplementation with nucleosides should rescue infection in KO lines. Addition of nucleosides to XRN1-KO cells failed to restore SINV infection (Fig.5c). However, ectopic nucleosides strongly improved viral fitness (∼7-fold) for the KO cells rescued with XRN1 WT, but not in KO cells rescued with the catalytically inactive XRN1 (Fig.5c). In agreement, nucleoside addition to XRN1 PKO moderately enhanced SINV infection (Fig.S5c). Addition of nucleosides to other lines with deleterious phenotypes, such as DCP2-KO and DDX6-KO also failed to rescue SINV infection (Fig.5d and S5d). By contrast, nucleotide supplementation rescued SINV infection in lines with less deleterious effects, such as DCP1-KO and PATL1-KO, reaching levels of viral gene expression similar to those observed in the parental line (Fig.5d). Altogether, these results suggest that nucleoside supplementation effectively overcomes 5-3DM derived defects in SINV replication, but only in conditions that are not fully deleterious for infection.

## DISCUSSION

XRN1 is a key regulator of cellular mRNA decay, but its role in infection is still under debate (Charley and Wilusz, 2016; Gaglia and Glaunsinger, 2010; Molleston and Cherry, 2017; Moon and Wilusz, 2013). Here, we propose a novel model in which XRN1 and the 5-3DM pathway can mediate the degradation of cellular RNAs in infected cells and supply free nucleotides for viral replication. This novel mechanism is well supported by the striking observation that the alphavirus RNA replaces the biomass of cellular mRNAs, reaching up to 70-80% (depending on the MOI) of the polyadenylated RNA pool (Fig.3c) (Garcia-Moreno et al., 2019). What triggers cellular mRNA degradation remains unknown; however, it is possible to speculate that protein synthesis shutoff caused by eIF2α phosphorylation could induce RNA decay (Buschauer et al., 2020; Carrasco et al., 2018; McInerney et al., 2005; Radhakrishnan et al., 2016; Radhakrishnan and Green, 2016; Weber and Chang, 2024). This could explain why cellular mRNAs that are translated in infected cells, such us interferons and ISGs, are not depleted after SINV infection (Li et al., 2015; Walsh et al., 2013).

We show that the exonuclease XRN1 is essential for the replication of several alphaviruses, and that the availability of a monophosphorylated 5’ for its activity is produced by the 5-3DM rather than the endonuclease activity of RNaseL (Fig.1 and S1, 2c and S2b) (Garcia-Moreno et al., 2019; Li et al., 2016). Additionally, we observed that XRN1 and the decapping complex accumulate in VROs (Fig.2d) and that the accumulation of these cellular proteins in replication sites correlates well with viral fitness (Fig.S2a). The connection between the decapping complex and XRN1 is underscored by the rescue experiment with an XRN1 mutant that cannot interact with the scaffold protein EDC4 and, consequently, cannot assemble a functional 5-3 DM (Brothers et al., 2023; Chang et al., 2019). In such conditions, XRN1 fails to restore SINV infection in XRN1 KO cells (Fig.2a), further supporting the relevance of the decapping machinery at inducing cellular RNA degradation in alphavirus infection.

It was proposed that the effect of XRN1-KO in infection could be due to an increased expression of ISGs. However, our RNAseq experiment revealed that most ISG-encoding mRNAs are not regulated in XRN1-KO cells, with a few exceptions that exhibit a subtle increase (Fig.S3d). However, the increase in the abundance of these RNAs does not robustly result in an increase on protein levels across KO clones with identical infection phenotypes (Fig.S3d-f) (BenDavid et al., 2022; Burgess and Mohr, 2015; Liu et al., 2021; Zou et al., 2024). This is also confirmed in 5-3DM KO cells which exhibits similar levels of expression of OAS1/3 and IFIT1 compared to WT (Fig.S3f). Moreover, other viruses that are susceptible to the antiviral programme exhibit no replication defects in XRN1-KO cells, including OC43, Influenza A virus (Fig.S1i-j), and HIV (Garcia-Moreno et al., 2019). Altogether, these findings support a mechanism for XRN1 function in SINV infection that does not involve the differential expression of ISGs.

Alternatively, similar to the well-studied role in flavivirus, XRN1 could be involved in the generation of sgRNAs via partial degradation of SINV genome. However, through the analysis of RNA sequencing and iCLIP2 in SINV infected cells, we did not identify an increase in read density across the viral genome in WT and XRN1 KO lines, downstream of the 3’ XRN1 binding peaks (Fig.S4f). Altogether, these findings highlight the difference between the XRN1-mediated infection in flavivirus and alphavirus.

SINV induces a pervasive degradation of housekeeping cellular mRNA that is replaced in biomass by viral RNA (Fig.3b-c) (Garcia-Moreno et al., 2019). However, it was unknown whether XRN1 was the mediator of the observed RNA decay. We applied iCLIP2 (Buchbender et al., 2019) to determine the scope of RNA in intimate contact with XRN1 in infected cells. Our results revealed a strong association at 4h post SINV infection and provided evidence of binding for nearly half of the degraded transcripts. This is an important overlap given that iCLIP2 suffers from low sensitivity due to the limitations of UV crosslinking (Esteban-Serna et al., 2023; Hafner et al., 2021; Lee and Ule, 2018) (Fig.4d-f). Our results thus reveal that XRN1 associates with a substantial proportion of the degraded RNAs in SINV infected cells.

Our data also reveals that XRN1 is tethered to the VROs in SINV infected cells while negligible binding to viral RNA is observed at 4hpi when cellular mRNA degradation begins (Fig.2d and 4g-i). The proximity of XRN1 (and plausibly cellular mRNA degradation) to the VROs may increase the local concentration of nucleotides. XRN1 nucleolytic reaction generates monophosphorylated nucleotides, which must be modified into tri-phosphorylated ones to be incorporated into nascent vRNA (Lane and Fan, 2015). Coherently, we also found the nucleotide modifying enzymes of the purine and pyrimidine salvage pathway in proximity to the VROs and their knockdown decreases SINV fitness (Fig.5a-b and S5a), being compatible with XRN1-mediated RNA degradation supplying with monophosphorylated nucleotides. In agreement, in other families of fast-replicating viruses such as picornavirus, the salvage pathway is required for viral infection (Nouwen et al., 2024). Finally, mimicking the XRN1-mediated release of nucleotides by providing external nucleosides, restores SINV replication in XRN1-WT-rescued cells as well as DCP1 and PATL1-KO cells, suggesting that nucleoside levels are a limiting factor for SINV, probably because of its high replication rate (Fig.5c-d). However, in XRN1-KO, XRN1 catalytically inactive mutant, DCP2-KO, and DDX6-KO the addition of nucleosides was not sufficient to restore infection (Fig.5c-d and S5d). The initial steps after virus entry are fundamental for the establishment of infection (Boersma et al., 2020). One possible explanation for the lack of SINV infection upon nucleosides addition in XRN1 and DCP2 KO cells is a highly penetrant defect for viral RNA to transition between translation and replication. In such scenario, nucleoside rescue might not be possible due to the lack of replication competent viral RNAs in the cell.

Taken together, our results support a model in which XRN1 is recruited to viral factories via interaction with nsP1, where it degrades cellular mRNAs, releasing mono-phosphorylated nucleotides. These are then converted into tri-phosphorylated nucleotides by the salvage pathway, resulting in a high local concentration sustaining the rapid replication rate imposed by SINV (Fig.S5e).

## ACKNOWLEDGEMENTS

A.C. is supported by a European Research Council (ERC) consolidator grant 101001634, MRC grants MR/R021562/1, MC_UU_12014/10, MC_UU_00034/2 and MC_UU_00034/5. V.R. was funded by the European Union’s Horizon 2020 research and innovation programme under Marie-Sklodowska-Curie n. 892756. R.H. is funded by the Swedish Foundations’ Starting Grant (Ragnar Söderberg Foundation), Swedish Research Council (Vetenskapsrådet), the Alex and Eva Wallström Foundation, and the Karolinska Institute Research Foundation. A.E.-B. is funded by Fundación Ramón Areces postdoctoral fellowship program. N.P. is funded by the MRC CVR PhD programme. N.R. is funded by the Marie Curie Individual Fellowship, RBP-ReguNet — HORIZON-MSCA-2021-DN-01. Z.R.D.L. was funded by WELLCOTR/WT_PHD_PROGRAMME 218518/Z/19/Z. S.M is supported by the EPSRC (V011359/1 (P)). V.P. is supported by an ERC synergy grant EPIC 101118521 and a Wallenberg Academy Fellowship KAW.2021.0167 and the Swedish Research Council VR 2022-05272. ID is funded by a Wellcome Investigator Award (209412) and by an NIH grant.

While this work is funded by the European Union, views and opinions expressed are however those of the author(s) only and do not necessarily reflect those of the European Union or the European Research Council Executive Agency. Neither the European Union nor the granting authority can be held responsible for them.

We thank the Castello, Fletcher, and Davis lab members for the scientific discussion over the development of this work. We thank Van Kuppeveldd for the CVB3-eGFP construct and anti 3A CVB3 antibody. We thank Alain Kohl and Andres Merits for the CHKV capsid antibody. We thank Sam Wilson for the OAS1/OAS3 antibody, RNaseL sgRNAs, and OC43 virus. We thank Juan Arriaza Garcia for VACV-mCherry. We thank Arvind Patel for anti DIII 1B ZIKV antibody. We thank Ed Hutchinson group for IAV virus.

## AUTHOR CONTRIBUTIONS

Conceptualization, V.R., I.D., A.C.; Methodology, V.R., L.I., R.H, N.P., J.H., A.C.; Investigation, V.R., L.I., R.H., A.E-B., S.N., N.P., N.R., Q.G., H.C., M.N., Z.R.D.L., J.J., M.N.; Writing, original draft, V.R and A.C.; Writing, editing, V.R., L.I., R.H., A.E-B., N.P., N.R., M.N., Z.R.D.L., C.I., D.L.R., D.L.R., J.H., V.P., I.D., A.C.; Funding acquisition, V.R., A.C.; Resources, C.I., D.L.R., V.P., S.H., I.D. A.C.; Supervision, V.R., I.D., A.C.

## MATERIALS AND METHODS

### Viruses and cells Cell culture

HEK293, HEK293T, and A549 cells were maintained in DMEM (Gibco, 41965039) with 10% foetal bovine serum (FBS) (Gibco, 10500064) and 1x penicillin/streptomycin (Sigma Aldrich, P4458) at 37**°**C with 5% CO_2_. BHK21 cells were used for producing virus stocks and were maintained in same conditions. HEK293-Flp-In T-REx (Thermo Fisher Scientific # R78007) were maintained in DMEM (Gibco, 41965039) with 10% foetal bovine serum (FBS) (Gibco, 10500064), 1x penicillin/streptomycin (Sigma Aldrich, P4458), 100 µg/ml Zeocin (Gibco, #R25001), and Blasticidin S Hydrochloride (Thermo Fisher Scientific, J67216.8EQ).

A549 and HEK293-Flp-In T-REx XRN1-KO cell lines were generated using TRANSIT-CRISPR (Sigma-Aldrich) as previously described (Garcia-Moreno et al., 2019).

HEK293-Flp-In T-REx XRN1-KO cells were co-transfected with pcDNA5/FRT/TO XRN1-WT-FLAG-HA, XRN1-D208A-FLAG-HA, or XRN1-Δ1650-1706-FLAG-HA plasmids and pOG44 (Thermo Fisher Scientific, #V600520) to generate add-back cell lines. The XRN1 protein expression was induced with doxycycline (1 µg/ml) treatment for at least 96 h. Cells were selected and maintained in DMEM (Gibco, 41965039) with 10% foetal bovine serum (FBS) (Gibco, 10500064), 1x penicillin/streptomycin (Sigma Aldrich, P4458), 150 µg/ml Hygromycin B (Invitrogen, #10687010), and 10 µg/ml Blasticidin S Hydrochloride at 37**°**C with 5% CO_2_. The plasmid expressing XRN1-D208A and Δ1650-1706 was generated by site-directed mutagenesis using pcDNA5-FRT-TO-XRN1-Flag-HA (Garcia-Moreno et al., 2019) as template. Primers are listed in Table 4. The mutagenesis was confirmed by sequencing.

HEK293T-DCP2, DCP1a/b, DDX6 (Weber and Chang, 2024), PATL1, EDC3, EDC4 knocked-out cell lines were generated with the CRISPR-Cas9 system (table 4). Guide RNAs used are listed in Table 4.

RNaseL knock-out cell lines were generated using CRISPR/Cas9 system via lentiviral transduction (Wickenhagen et al., 2021) in both A549 and HEK293 cell lines.

ACE2 expressing cell lines (HEK293T, WT, DCP2, DCP1a/b, DDX6, PATL1, EDC3, EDC4 - KO and A549-WT, Partial XRN1-KO, and Full-KO) were generated by transducing lentiviral particles obtained from co-transfecting the plasmids pLV-hygro-ACE2 (3 µg), VSV-G (3 µg), and nLPG (1 µg) into HEK293T using lipofectamine 3000 (Invitrogen, L3000001) following manufacturer’s instructions.

### Viruses

Sindbis virus was produced by *in vitro* transcription from the plasmid pT7-SVwt (Castello et al., 2006). SINV_-nsP3-mScarlet_ was generated by inserting the mScarlet sequence into pT7-SVwt plasmid between the SpeI restriction site (Kamel et al., 2024). Replicative alphaviruses with fluorescent reporters (i.e. SFV-mCherry, RRV-mCherry, and ONNV-mCherry) were generated by inserting mCherry after a duplicated subgenomic promoter as previously described for SINV-mCherry (Garcia-Moreno et al., 2019). Chikungunya virus (CHKV) is part of the Eastern/Central/South African (ECSA) genotype, ICRES-1 strain. Coxsackie B3 virus (CVB3) has eGFP sequence followed by a 3Cpro cleavage site inserted between the 5’UTR and the coding sequence (Lanke et al., 2009). Vaccinia virus is mCherry tagged and is a gift from Juan Arriaza Garcia. Influenza A virus/Puerto Rico/8/1934 (H1N1, PR8) is a gift from the Ed Hutchinson lab. ZIKV-WT MR766 (ATCC VR-84 strain). ZIKV-mCherry is a gift from Andres Merits lab and are described in (Mutso et al., 2017). HCoV-OC43 strain MA-P9 (ATCC VR-1558) is a gift from Sam Wilson lab. For SARS-CoV-2 infection experiments we used the isolate CVR-GLA-1 strain (Rihn et al., 2021).

### Cell viability

Equal number of cells were seeded into a 12 wells plate and counted at the indicated days post seeding. Total number of cells and percentage of live cells was assessed by trypan blue staining and automatically counted with the Countess II FL automated Cell Counter (Thermo Fisher Scientific).

### siRNA transfection

1.0×10^4^ A549 cells were seeded in 24 well plate and transfected with a mix of siRNAs (Silencer select, validated) to a total amount of 25 pmol (table 4) using Lipofectamine RNAimax (Invitrogen, 13778075) following the manufacturer’s instructions. 48 hours post transfection, cells were infected with SINV_-nsP3-mScarlet_ (MOI 0.1) in serum-free media for 1 h, then media was replaced with 5% FBS. 18hpi cells were washed with PBS and lysed in RIPA buffer (0.1% SDS, 0.5% sodium deoxycholate, 150 mM NaCl, and 50mM HEPES) for western blot analysis.

### Nucleoside complementation experiments

0.2×10^6^ cells (HEK293 and HEK293T) were seeded in 24 well plate in presence or absence of the nucleoside mix solution (EmbryoMax Nucleoside 100x, Millipore EMD #ES-008-D) at 2x final concentration. Next day, cells were infected with SINV-WT (MOI 0.1), lysed, and collected for western blot analysis 18 hours post infection.

### Cell lysis and Western blot

To prepare whole cell lysate for Western blotting analysis, cells were washed with PBS and resuspended in RIPA buffer (0.1% SDS, 0.5% sodium deoxycholate, 150 mM NaCl, and 50mM HEPES, pH 7.9 and 0.1 mM AEBSF serine protease inhibitor) supplemented with Benzonase (1 µl/ml) and incubated on ice for 30 min. Next, total lysate was centrifuged at max speed for 10 min. Samples were resolved on SDS-PAGE and analysed by Western blotting using specific antibodies. Li-Cor Odyssey system for visualization and Image Studio Lite software (Li-Cor) for quantification. Antibodies used are listed in Table 4.

### Protein-protein interaction analysis

6×10^6^ HEK293 Flp-In T-REx XRN1-eGFP (XRN1-GFP) or HEK293 Flp-In T-REx eGFP-Ctrl (GFP-Ctrl) cells were seeded in a 10 cm dish in DMEM supplemented with 10% FBS. XRN1-GFP cells were induced with 1 μg/ml of doxycycline overnight, while GFP-Ctrl cells were induced 6 h prior infection. Cells were infected with 2 MOI of SINV-WT for 18 hours. After harvesting, cells were washed twice with ice-cold PBS and lysed in 1 ml of lysis buffer (10 mM Tris/Cl pH 7.5, 150 mM NaCl, 0.5 mM EDTA, 0.5 % Nonidet™ P40 with 0.1mM AEBSF serine protease inhibitor) for 30 min on ice. Samples were centrifuged for 10 min at 4 °C at 17,000xg. The cleared supernatant was then incubated for 3 h at 4 °C with shaking with 25 μl ChromoTek GFP-Trap® Magnetic Agarose beads (Proteintech) previously equilibrated in lysis buffer. After three washing steps (10 mM Tris/Cl pH 7.5, 150 mM NaCl, 0.05 % Nonidet™ P40 substitute and 0.5 mM EDTA), beads were incubated with wash buffer supplemented with Benzonase (1U/mL) for 15 min at 37°C and washed again twice with wash buffer without Benzonase. To elute, beads were resuspended in 50 μL 1% SDS and incubated for 5 minutes at 55°C with rotation (1100rpm).

### Protein-protein interaction analysis for mass spectrometry

5×10^6^ HEK293 Flp-In T-REx expressing eGFP or XRN1-eGFP were induced with 1 µg/mL doxycycline for 8 or 96 h, respectively. Cells were infected with 10 MOI of SINV or mock infected in FBS-free DMEM and incubated for 1 h. Media was replaced with 5% FBS-containing DMEM and incubated for 18 h. Cells were lysed in 1 mL lysis buffer (10 mM Tris/HCl pH 7.5, 150 mM NaCl, 0.5 mM EDTA, 0.5 % Nonidet™ P40 Substitute, 0.1 mM AEBSF serine protease inhibitor) containing 1 µL/mL Benzonase (Millipore, #70746-4). 25 µL of GFP-Trap agarose beads were equilibrated in lysis buffer and incubated with the whole cell lysate for 2 h at 4°C with gentle rotation. Beads were washed five times with 500 µL lysis buffer. An additional wash step was performed with 500 µL of wash buffer (10 mM Tris/HCl pH 7.5, 150 mM NaCl, 0.5 mM EDTA, 0.1 mM AEBSF serine protease inhibitor) and proteins were eluted twice with 50 µL glycine-elution buffer (200 mM glycine pH 2.5) and neutralized with 15 µL neutralization buffer (1 M Tris pH 10.4). Three biological replicates were used for the proteomic experiment.

### Mass spectrometry

XRN1 IP eluates were cleared and digested with single-pot solid-phase-enhanced sample preparation (SP3) protocol using carboxylate modified magnetic beads (Sigma-Aldrich) (Hughes et al., 2014; Kamel et al., 2021). Samples were denatured in 200 µl of Urea buffer (8 M urea, 100 mM ammonium bicarbonate in water), and subjected to reduction and alkylation in one solution environment by addition of 10 mM Tris[2-carboxyethyl] phosphine hydrochloride and 50 mM 2-chloroacetamide and 30 min in-dark incubation. Protein binding was performed at 77% acetonitrile, digestion was performed on beads with MS-grade trypsin protease (Promega) after 10 cycles of washes. Digestion products were acidified with formic acid (final concentration 5%), then desalted with C18 tip packed in-house with C18 resin (SPE-Disks-C18, Affinisep) prior to LC-MS/MS analysis. Desalting eluates were dried using SpeedVac, and reconstituted with MS loading buffer (5 % DMSO, 5 % formic acid in water) before LC-MS/MS analysis.

Peptides were separated by nano liquid chromatography (Thermo Scientific Ultimate RSLC 3000) coupled in line a Q Exactive mass spectrometer equipped with an Easy-Spray source (Thermo Fischer Scientific). Peptides were trapped onto a C18 PepMac100 precolumn (300 µm i.d. x5 mm, 100 Å, Thermo Fischer Scientific) using Solvent A (0.1% Formic acid, HPLC grade water). The peptides were further separated onto an Easy-Spray RSLC C18 column (75 μm i.d., 50 cm length, Thermo Fischer Scientific) using a 60 minute linear gradient (15% to 35% solvent B (0.1% formic acid in acetonitrile)) at a flow rate 200 nl/min. The raw data were acquired on the mass spectrometer in a data-dependent acquisition mode (DDA). Full-scan MS spectra were acquired in the Orbitrap (Scan range 350-1500m/z, resolution 70,000; AGC target, 3e6, maximum injection time, 50 ms). The 10 most intense peaks were selected for higher-energy collision dissociation (HCD) fragmentation at 30% of normalized collision energy. HCD spectra were acquired in the Orbitrap at resolution 17,500, AGC target 5e4, maximum injection time 120 ms with fixed mass at 180 m/z. Charge exclusion was selected for unassigned and 1+ ions. The dynamic exclusion was set to 20 s.

Protein identification and quantification were performed using Andromeda search engine implemented in MaxQuant (2.0.1.0). Mass spectra were searched against reference proteome datasets: human proteome (Uniprot_id: UP000005640, downloaded Nov 2016) and a custom database including all the known SINV polypeptides. The following search parameters were used: full tryptic specificity with maximal two missed cleavage sites, carbamidomethyl (C) set as fixed modification, acetylation (protein N-term) and oxidation (M) set as variable modifications. Match between runs was turned on, samples from the same experimental condition were assigned as neighbouring fractions. False discovery rate (FDR) cut-off for peptide identification was set to 1%. All other settings were set to default.

### Proteomic quantitative analysis

The proteinGroup file of MaxQuant search result was imported in RStudio (R Project) for further processing. Proteins flagged as potential contaminants, reverse, and only identified by sites were removed. Proteins that have less than 2 valid intensity measurements in all conditions were removed prior to downstream analysis. XRN1 interactome was determined through contrasts of XRN1-IP versus GFP-IP in either SINV-infected or mock-infected cells. Normalisation was performed to samples in each condition using variance stabilisation normalisation (vsn) method (Huber et al., 2002). Imputation of missing values was only preformed for proteins that have missing values in all replicates in one experimental condition (“absent”), while present in the other condition. Only missing values in the “absent” condition were imputed using deterministic minimum method with package ImputeLCMD (Lazar et al., 2016). Statistical testing was performed based on empirical Bayesian model moderated T-test using package limma (Ritchie et al., 2015). P values obtained by moderated T-test were adjusted using Benjamini-Hochburg method. Proteins with FDR < 0.05 and log2 fold change > 0 were classed as XRN1 interactors.

For analysis of the contrast between XRN1 interactors in SINV- and mock-infected cells, only proteins determined as XRN1 interactors in previous analysis were included. Normalisation was performed for all samples using vsn method, missing value imputation was only performed on the “absent” group for proteins that exhibit “present-absent” pattern, and statistical testing was performed using moderated T-test in R package limma. For analysis of contrasts between XRN1 interactors and whole cell proteome (WCP), the WCP result were obtained from previous publication (Garcia-Moreno et al., 2019). Normalisation, missing value imputation, and statistical analysis was performed using the same method as in analysis of contrast between XRN1 interactors in SINV- and mock-infected cells.

### Immunofluorescence and smFISH

smFISH experiments were performed as described in (Lee et al., 2022) with few modifications. 5.0×10^4^ A549 or 1.5×10^5^ HEK293 cells were seeded on pre-washed sterile High Precision Coverslips (Merienfeld, # 0117520) in a 24well plate and incubated in DMEM with 10% FBS. Cells were either mock-infected or infected with SINV at specific MOI indicated in figure legends. Cells were fixed at indicated time points with 4% formaldehyde for 15 min at room temperature, washed with 1x PBS and permeabilised with 0.1% Triton X-100 in PBS for 10 min at room temperature. Cells were washed twice with 1x PBS and 2X SSC, followed two pre-hybridization steps (20 min each at 37°C in pre-warmed wash solution (2× SSC, 10% formamide). Hybridisation was performed over-night at 37°C in hybridization solution (2× SSC, 10% formamide, 10% dextran sulphate) supplemented with SINV RNA-specific Stellaris (LGC Biosearch Technologies) and oligo(dT)25 (Life technologies Ltd) probes (1:100 of 25 μM stock probe). From this point on cells were kept in the dark). Cells were then washed twice for 20min in wash buffer at 37°C, followed by incubation with DAPI (1 µg/ml) in wash solution at room temperature for 10 min. Finally, cells were washed twice in 2X SSC buffer for 5min. At this step cells are either dipped in pure water and mounted on ProLong Diamond antifade mounting media (Invitrogen) or washed twice with 1X PBS to continue with immunofluorescence. For the latter, the PBS washes were followed by the addition of blocking solution (5% BSA, 0.1% Tween 20, in 1x PBS) for 1h at room temperature. This was subsequently replaced by a blocking solution supplemented with the primary antibody (table 4) for 1h at room temperature, followed by three washes in 0.1% Tween-20 1x PBS. Blocking solution supplemented with AlexaFluor-fluorescently conjugated secondary antibodies was then added to the cells for 1h at room temperature, followed by three washes in 0.1% Tween-20 1X PBS and once in 1X PBS. Finally, coverslips were dipped in pure water and mounted using ProLong Diamond antifade mounting media (Invitrogen).

When performing immunofluorescence alone, permeabilised cells (0.1% Triton X-100 in 1x PBS) were washed three times in 1x PBS followed by the addition of blocking solution (5% BSA, 0.1% Tween 20, in 1x PBS). The same protocol used for smFISH + immunofluorescence (described above) was then followed.

Images were acquired on a Zeiss LSM880, using a 63x oil Plan-Apochromat DIC M27 objective (NA 1.4). Mean average signal for at least 100 cells from three independent experiments was measured with Fiji. smFISH and Immunofluorescence images were prepared using Zeiss Zen 3.4 Blue Edition and Fiji software. Line profiles were generated using Fiji “plot-profile” tool.

### RNA extraction and RT-qPCR

Cells were harvested at indicated time points and centrifuged for 5 min at 1000 rpm. Cell pellets were washed once with PBS and resuspended in TRIzol Reagent (ThermoFisher scientific). RNA was extracted with chloroform and precipitated in isopropanol according to the manufacture instructions. RNA pellets were washed twice with 80% ethanol and suspended in RNAse-free water. Reverse transcription and RT-qPCR analysis was performed by Luna universal one-step RT-qPCR kit (NEB # E3005L) with the primers listed in Table 4.

### iCLIP2 experiments

HEK293 Flp-In T-REx XRN1-GFP (Garcia-Moreno et al., 2019) were induced with 1 µg/ml of Doxycycline for 96 hours. Cells were infected with SINV-WT (10 MOI) for 4 and 18 hpi. iCLIP2 experiments were performed as described in (Garcia-Moreno et al., 2023). XRN1-IP libraries and SMI libraries were each individually pooled equimolarly and then mixed at the following proportions: 57% XRN1-IP pool and 43% SMI pool. The pooled pool was sequenced on a NextSeq 550 sequencer with a 75 cycle High-output cartridge (Illumina, #20024906).

### iCLIP2 analysis

Raw FASTQ files were demultiplexed using the Je Suite (Girardot et al., 2016) and adapters were trimmed using Cutadapt (Martin, 2011). STAR was used to align reads to a concatenated human (GRCh38, ENSEMBL Release 106) and SINV (pT7-SVwt) genome in end-to-end alignment model (Dobin et al., 2013). Only uniquely aligned reads were retained for downstream analysis. PCR duplicates were collapsed using unique molecular identifiers (UMIs) with the Je Suite. The crosslink truncation site for each read (-1 from the 5’ start site of the read) was extracted using BED Tools (Quinlan and Hall, 2010). Peak calling was performed with HTSeq-clip and the R/Bioconductor package, DEW-seq (Sahadevan et al., 2022). HTSeq-clip was used to generate a sliding window annotation of the human and SINV genome (50nt window, 20nt step size) and calculate the frequency of crosslink truncation sites within each window. DEW-Seq was then used to calculate the differential enrichment of each window relative to size matched input control samples, with a cut-off of >2 log2 fold change and <0.01 adjusted p-value. Multiple hypothesis correction was performed using the Independent Hypothesis Weighting (IHW) method (Ignatiadis et al., 2016). Overlapping windows were merged to form binding regions. PCA analysis was performed using DESeq2. Following size correction and variance stabilisation, the 1000 most variable sliding windows were selected and used for PCA plotting. Binding site properties, including gene name, biotype, gene feature, were extracted from the ENSEMBL genome annotation using the Genomic Ranges package in R. Metagene analyses were performed using functions from the cliProfiler package in R. SINV genome coverage in reads per million was calculated using BED Tools. SMI signal was subtracted from IP signal for plotting.

### RNA sequencing experiment

0.9×10^6 HEK293 WT, XRN1 Partial-KO, and XRN1-KO cell lines (Garcia-Moreno et al., 2019) were seeded in 6 well plates and infected with SINV (1 MOI) for 8 and 18 hpi. At the indicated time point, cells were resuspended in 1ml media; 200 µl of cells suspension was taken for western blot experiment and lysed in protein sample buffer, while the remaining 800 µl were resuspended in 1 ml Trizol for RNA extraction. The samples were stored at -80 °C immediately after collection and subsequently RNA was extracted with phenol/chloroform as described above.

Samples underwent an RNAClean XP bead (Beckman) cleanup to remove phenol residue and RNA quality was checked by Qubit Fluometer 4 (Thermofisher) and TapeStation 4000 (Agilent). Using the Illumina Stranded Total RNA Prep, Ligation with Ribo-Zero Plus kit (#20040529), 100ng of total RNA was taken for library preparation. Following manufacturer guidelines, RNA was subjected to probe based ribosomal RNA depletion followed by RNA fragmentation, reverse transcription and then converted to dsDNA. After end repair, A tailing, and ligation of specific adaptors, samples were PCR amplified with indexing primers. Libraries were pooled in equimolar concentrations and sequenced in Illumina NextSeq 550 sequencer with a 75 cycle High-output cartridge (Illumina, #20024906). At least 85% of the reads generated presented a quality score of 30 or above.

### RNA sequencing analysis

T3’-sequencing adapter trimming and quality filtering were applied using Cutadapt V4.0 (Martin, 2011), with the parameters --nextseq-trim=20 and --minimum-length 28. Reads were mapped using Star V2.7.1a (Dobin et al., 2013) to the combined human genome (hg38) with SINV sequence as the reference. Read coverage per gene was counted using Subread (featureCounts) (Liao et al., 2014). Global transcription is shut off during SINV infection (Gorchakov et al., 2005), which may underestimate differential expression results if normalisation is carried out assuming unchanged RNA abundances. Appropriate normalisation genes were selected by Two One-sided t-test (TOST) analyses with unequal variances for significantly non-changing genes. 11 genes were identified as significantly unaffected by viral infection (Mock vs. 18 hpi, p < 0.1, d < 1.5 SDs) (Garcia-Moreno et al., 2019), and XRN1 knockout (this study, Mock WT vs. Mock XRN1 KO, p < 0.1, d < 1.5 SDs). Samples with less than 10 million read counts were removed and genes with less than 50 counts across all conditions were excluded from the analysis. Differential gene expression analysis was performed using DESeq2 (Love et al., 2014) with the parameter cooksCutoff=FALSE. The threshold for differentially expressed genes was defined as adjusted p-value < 0.05 and |log2(fold-change)| > 0.585.

### Codon optimised Luciferase reporter

The pcDNA5-Rluc-WT, up, or down constructs containing the wild-type DNA Renilla luciferase sequence from pRL-CMV (Promega AF025843) or the di-nucleotide optimized Rluc sequence optimised to resemble SINV (Upregulated transcripts, up) or Mock (downregulated transcripts, down) conditions were purchased from GeneScript and inserted into pcDNA™/FRT/TO (ThermoFisher # V652020) via BamHI and NotI sites. These sequences were amplified using primers containing the T7 sequence (table 4). Transcription and poly(A) tailing reactions were performed with these PCR products using mMESSAGE mMACHINE™ T7 Transcription (ThermoFisher #M0531S) and Poly(A) Tailing kit (ThermoFisher #AM1350) according to manufacturer’s instructions. RNA was extracted using RNeasy kit (Qiagen).

1µg of the WT, up, or down Rluc RNAs (modified di-nucleotide usage) were transfected into cells 2 h post SINV_-Nsp3-mScarlet_ infection using 1 MOI of virus in a 12-well plate. Cells were collected 4- and 6-hours post-transfection (hpt) for Luciferase measurement using Dual-Luciferase® Reporter Assay System (Promega) and RNA extraction for RT-qPCR assay using Luna® Universal One-Step RT-qPCR (New England Biolabs) and primers listed in table 4.

### Feature selection of codon usage

The usage of the existing 64 codons, defined as features, was calculated in each nucleotide sequence across the human genome. The values of codon usage are available from this previous study (Chai et al., 2022). Apart from codon usage, synonymous dinucleotide usage (SDUc) was also used as a feature. In this study, the SDUc of coding sequence across the human genome was calculated via the package DinuQ (Lytras and Hughes, 2020). We also used DinuQ to calculate the synonymous dinucleotide usage of the Sindbis viral genome (NCBI Reference Sequence: NC_001547.1).

We calculated the feature importance for the classification of different groups (the up-regulated vs down-regulated differential expressed genes of mock vs 18h) based on the gradient boosting machine (GBM) model from the feature selection package Caret (Kuhn, 2008).

---

**Figure S1.**
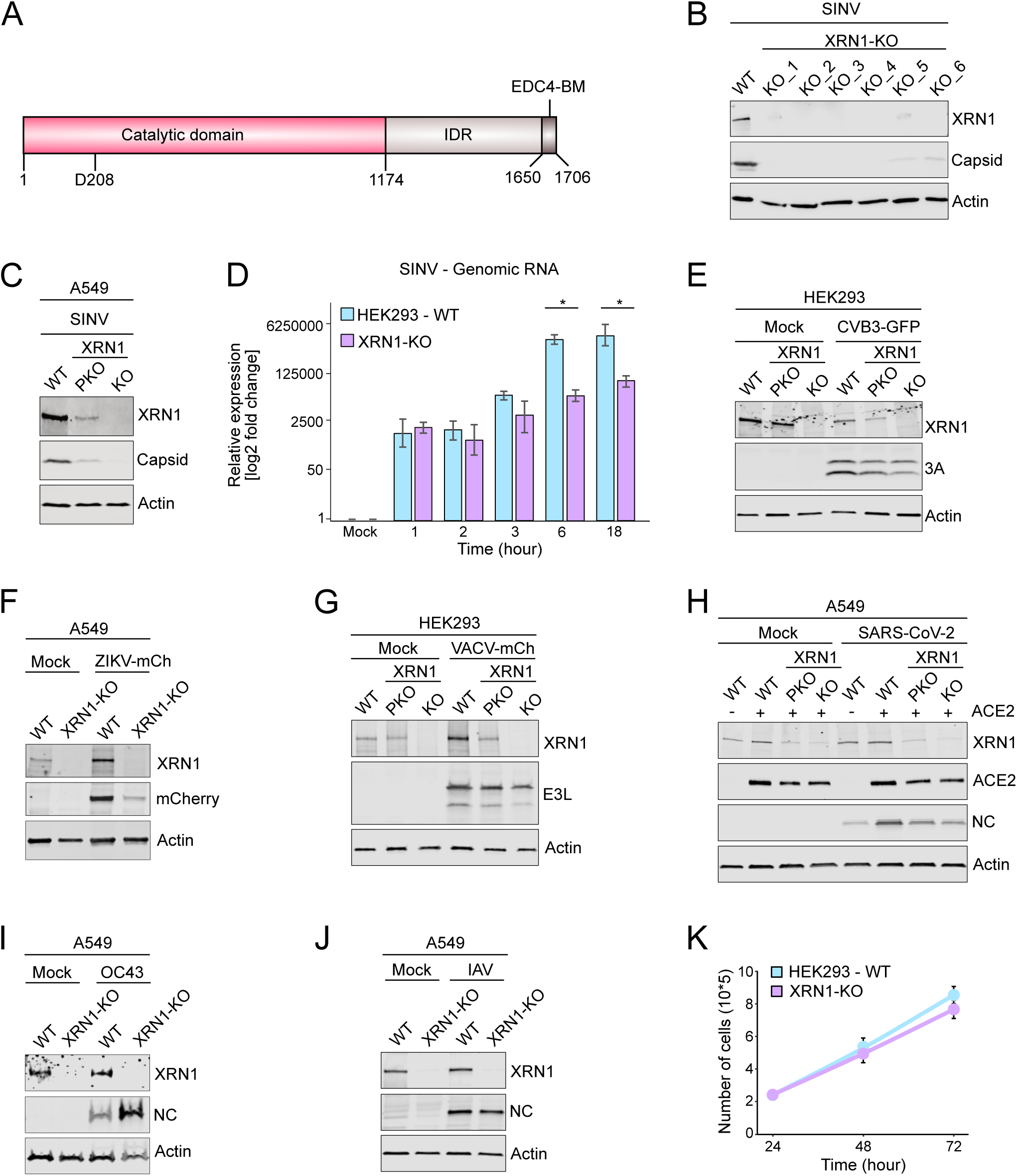
XRN1 influences the infection of different viral families. (A) Schematic representation of XRN1’s main features with amino acid positions. IDR: intrinsic disordered region; EDC4-BM: EDC4-binding motif; D208: catalytic residue. (B) Western blot analysis of SINV infection in HEK293-Flp-In T-REx WT and different XRN1 KO clones (KO_1-KO_6) (MOI = 0.5). n = 3. (C) Western blot analysis of A549-WT, XRN1 partial-KO (PKO), and full KO (KO) cells infected with SINV for 18 h (MOI = 0.1). n = 3. (D) RT-qPCR analysis of SINV genomic RNA in WT and XRN1-KO cells infected with SINV for the indicated time (MOI = 3). n = 3; error bars: standard error; * p < 0.1. (E-J) Western blot analysis of WT, XRN1 partial-KO (PKO), and full KO (KO) cells infected with CVB3-GFP (18 h. MOI = 0.1) (E), ZIKV-mCherry (mCh) (24 h. MOI = 0.5) (F), VACV-mCherry (18 h. MOI = 0.1) (G), SARS-CoV-2 (24 h. MOI 0.1) (H), OC43 (96 h. MOI = 0.1) (I), Influenza A virus (IAV, 24 h, MOI = 0.5) (J). n = 3. (K) Growth curve of WT and XRN1-KO cells. n = 3; error bars: standard error.

**Figure S2.**
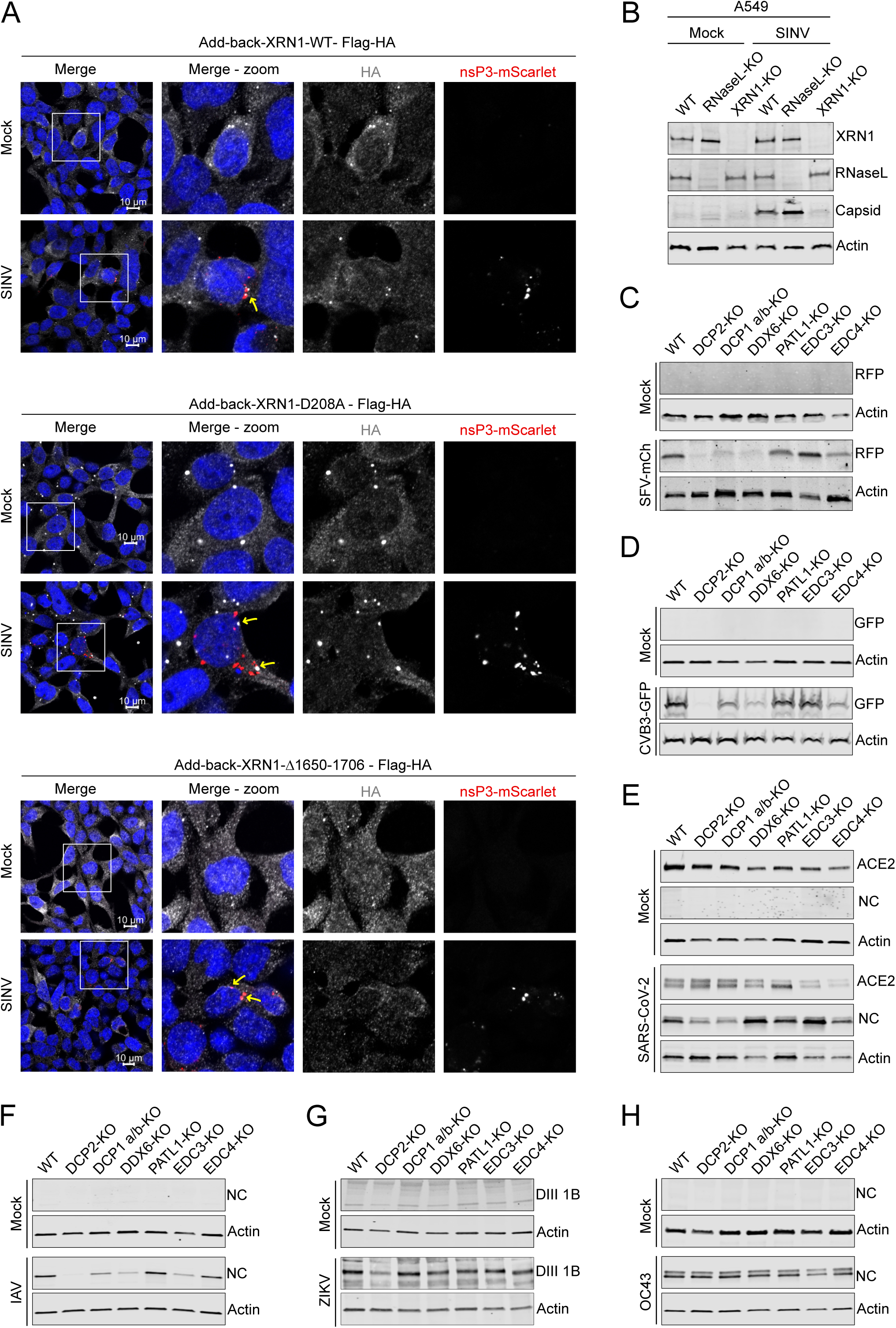
5’-3’ RNA degradation machinery influences the infection of different viral families. (A) Fluorescence microscopy analysis of XRN1-KO cells complemented with the indicated XRN1 mutant induced by doxycycline treatment 96 h before SINV infection for 18 h (MOI = 3, n = 2). XRN1 mutant protein was visualized by immunofluorescence with anti-HA antibody, nsP3 localisation was determined by detection of scarlet tag, and nuclei were stained with DAPI. Yellow arrows indicate colocalization of XRN1 and nsP3. (B) Western blot analysis of A549 WT, RNaseL-KO, and XRN1-KO cells infected with SINV for 18 h (MOI = 0.1). n = 3. (C-H) Western blot analysis of the indicated RNA decay KO cells infected with SFV-mCherry (18 h. MOI = 0.01) (C), CVB3-GFP (18 h. MOI = 0.1) (D), SARS-CoV-2 (24 h. MOI = 0.01) (E), IAV (24 h. MOI = 0.1) (F), ZIKV (24 h. MOI = 0.1) (G), and OC43 (72 h. MOI = 0.1) (H). n = 3.

**Figure S3.**
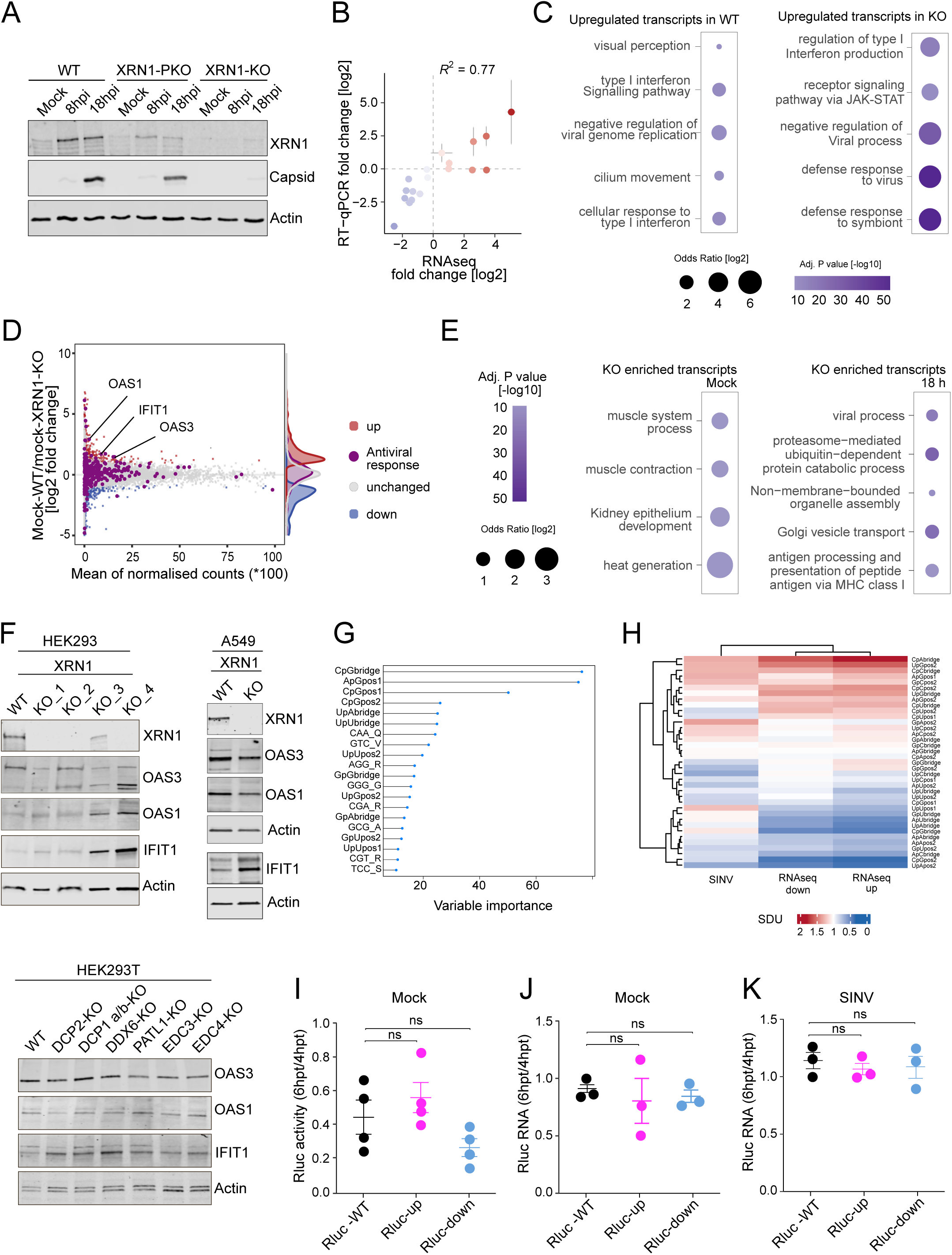
XRN1-KO transcriptome is comparable to WT cells. (A) Western blot analysis of samples from figure 3B (MOI = 1. n = 3). (B) Correlation of the RNAseq and RT-qPCR data from (Garcia-Moreno et al., 2019) for randomly selected transcripts. Error bars represent standard error of three independent experiments. (C) GO term enrichment for upregulated transcripts in SINV infected WT (left) and XRN1-KO (right) cells. (D) MA plot comparing the read coverage and the log2 fold change between WT and XRN1-KO uninfected cells of each gene detected in the RNAseq experiment from figure 3B. Red, blue, and grey dots represent significantly upregulated, downregulated, and unchanged transcripts levels, respectively. Purple dots represent ISGs in HEK293 cells, as defined in (Chen et al., 2024). Density plots displaying distribution of fold changes among each colour group are shown at the right side of the figure. (E) GO term enrichment of transcripts upregulated in XRN1-KO cells in non-infected conditions (XRN1 WT versus KO cells in non-infected conditions, left) and transcripts upregulated in XRN1-KO cells 18 hpi (XRN1 WT versus KO cells in infected conditions, right). (F) Western blot analysis of antiviral response proteins OAS1, OAS3, and IFIT1 in WT and XRN1/5’-3’ RNA decay components KO cells in non-infected conditions. n = 3 (G) Top 20 most important features and their corresponding importance scores identified by Synonymous Dinucleotide Usage (SDU) and Codon Usage comparing up versus down-regulated transcripts upon SINV infection (RNAseq experiment fig 3B). (H) Comparison of the SDU between SINV genome and cellular transcripts upregulated and downregulated upon SINV infection. (I) Luciferase activity of Rluc-WT or reporters with altered dinucleotide codon usage (Rluc-up and Rluc-down) measured at 6 hours post transfection (hpt) relative to the luciferase levels at 4 hours post transfection (hpt) in mock conditions. n = 4; error bars: standard error. (J-K) RNA fold change of RLuc-WT or reporters with altered dinucleotide codon usage (RLuc-up and RLuc-down) measured at 6 hours post transfection (hpt) relative to RNA levels at 4 hours post transfection (hpt) in mock (J) and infected conditions (K). n = 3; error bars: standard error.

**Figure S4.**
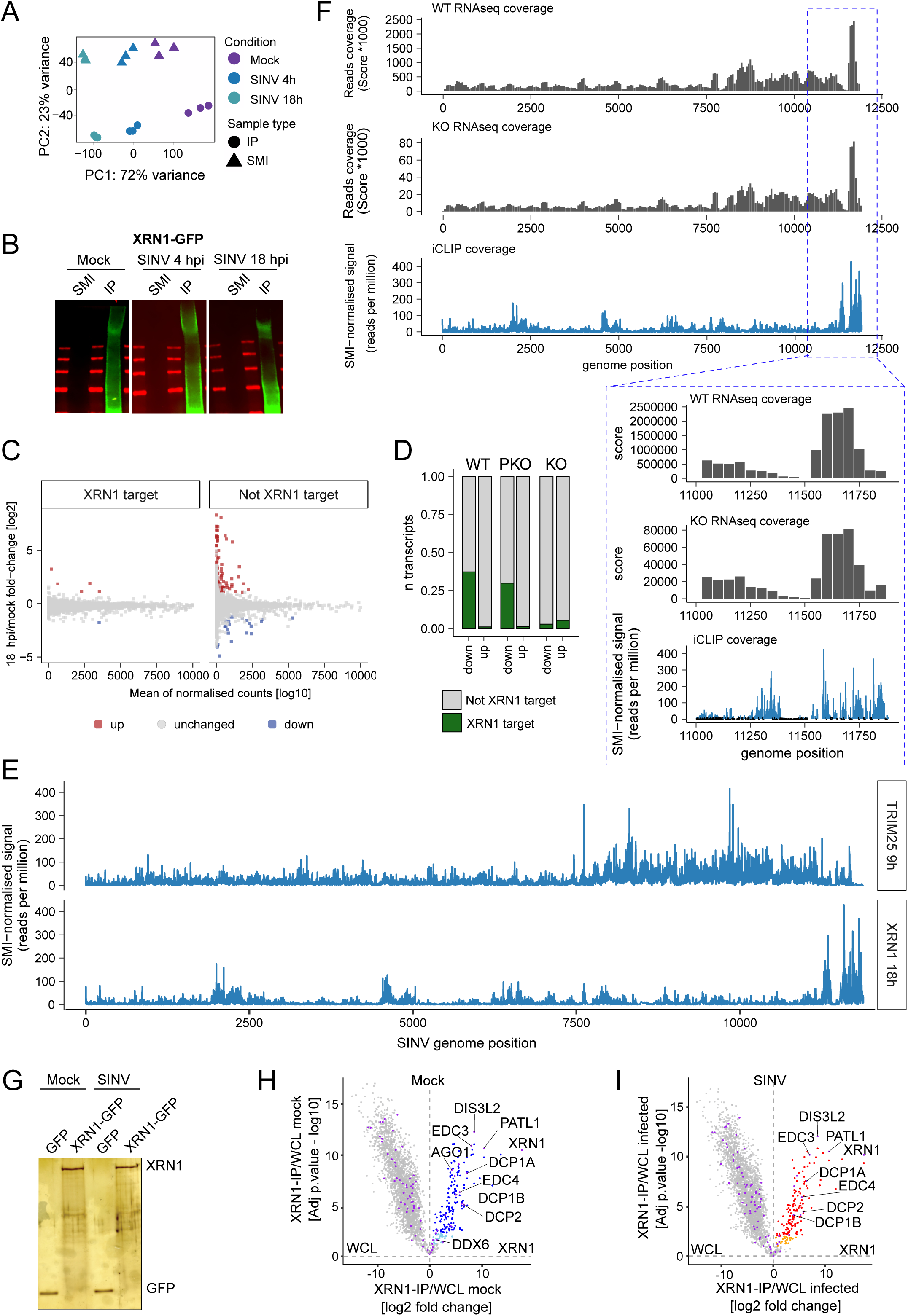
XRN1’s interaction profiles are consistent with its role in RNA degradation. (A) Principal component analysis of data from immunoprecipitated (IP) and size match input (SMI) samples from iCLIP2 experiments in uninfected and SINV-infected cells at 4 or 18 hpi. (B) Visualisation of IR adaptor of size match input (SMI) and immunoprecipitated (IP) samples from iCLIP2 experiment, in mock and infected conditions. (C) MA plot comparing the read coverage and the log2 fold change between the differentially regulated transcripts in XRN1-KO cells pre- and post-infection with the XRN1 targets (left) and not targets (right) identified by iCLIP2. Blue and red dots represent significantly downregulated and upregulated RNAs, respectively. (D) Bar plots reflecting the same data shown in figure 4F. Instead of absolute counts, the percentage of transcripts is plotted to better visualize the effect seen in KO cells. (E) Comparison of binding profile of TRIM25 (Álvarez et al., 2024) and XRN1on SINV genome at 9 and 18hpi, respectively. (F) SINV genome coverage from RNAseq (WT and XRN1-KO samples) and iCLIP2 data with inset showing a ‘zoomed in’ view of the 3’ end of the viral genome. (G) Silver staining analysis of samples from anti-GFP-beads for mass spectrometry analysis from figure 4 H-I. (H-I) Volcano plots showing XRN1 interactor proteins enriched in XRN1-IP compared to WCL in mock (H) and infected (I) conditions.

**Figure S5.**
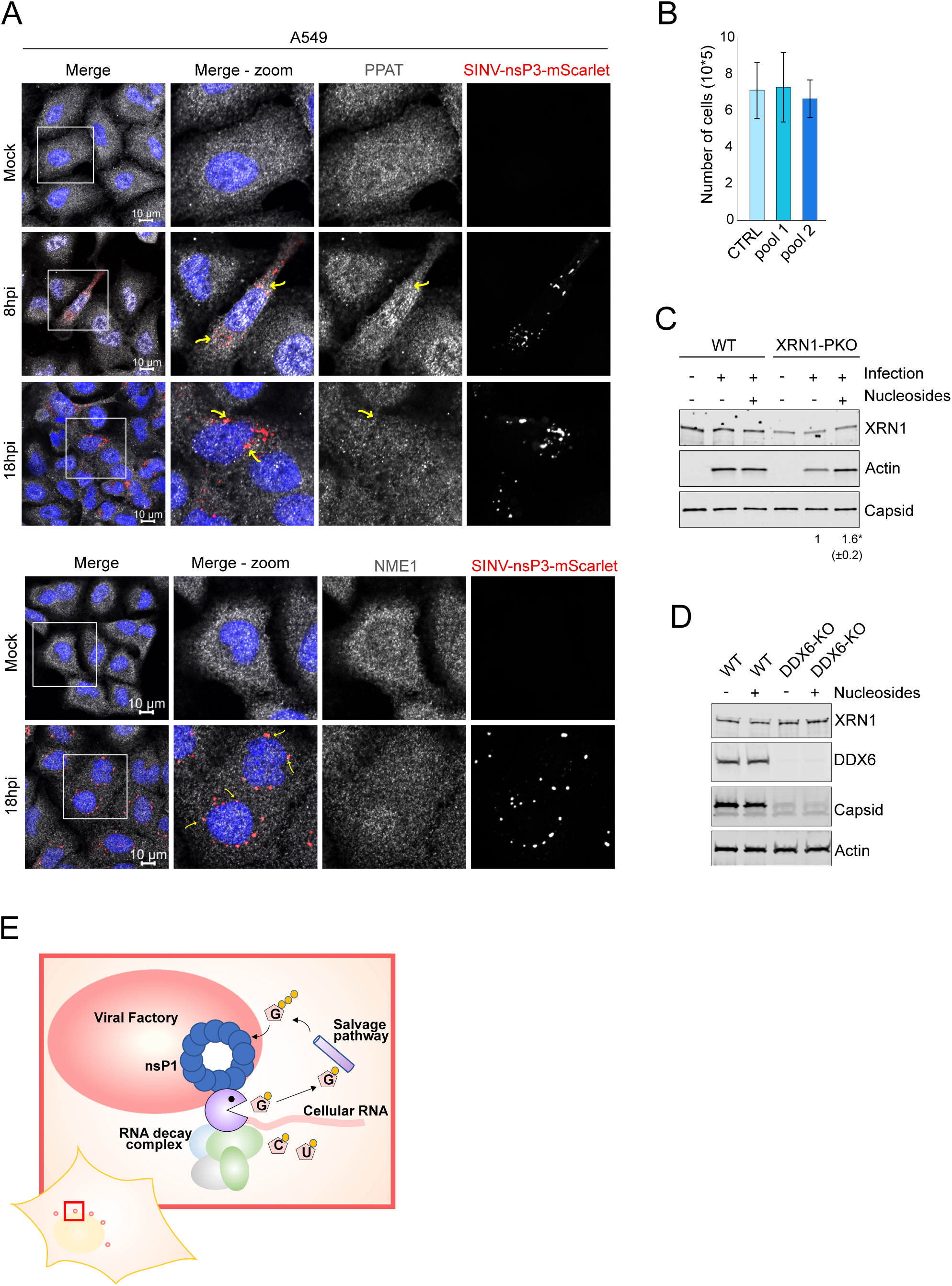
Enzymes involved in wider nucleotide metabolism are also proximal to VROs. (A) Fluorescence microscopy analysis of mock and SINV-infected cells at the indicated time-points post infection (MOI 1). n = 3. PPAT or NME1 were detected by immunofluorescence, SINV-nsP3 was visualized by tagging with mScarlet, and nuclei were stained with DAPI. Yellow arrows indicate colocalization of nsP3 with PPAT (top), or NME1 (bottom). (B) Cell count of WT cells and cells depleted of components of the nucleotide salvage pathway with two different siRNA mixes (Pool-1: HPRT1, CTPS1/2, APRT, UPRT, CDA, and NME3; and Pool-2: HPRT1, CTPS1, APRT, CDA, and UPRT), infected with SINV for 18 h (MOI 0.1). n = 3. Error bars: standard error. (C) Western blot analysis of HEK293 WT and XRN1 partial KO (XRN1-PKO) cells supplemented with nucleosides and infected with SINV for 18 h. (MOI 0.1). n = 5. Below the blots the average normalized capsid level of four independent experiments, in bracket standard error. * p < 0.05. (D) Western blot analysis of DDX6-KO cells, supplemented with nucleosides and infected with SINV for 18 h. (MOI 0.01). n = 3. (E) Schematic model representing the role of the RNA decay machinery and nucleotide salvage pathway in regulating alphavirus infection. XRN1 and the 5-3DM localize to SINV viral factories and degrade cellular transcripts in proximity to the replication centre to provide high local concentration of nucleotides to sustain viral replication.

